# Engineered Ubiquitin Variants Mitigate Pathogenic Bacterial Ubiquitin Ligase Function

**DOI:** 10.1101/2024.05.01.592009

**Authors:** Bradley E. Dubrule, Ashley Wagner, Wei Zhang, Adam J. Middleton, Adithya S. Subramanian, Gary Eitzen, Sachdev S. Sidhu, Amit P. Bhavsar

## Abstract

During infection some pathogenic gram-negative bacteria, such as *Salmonella*, manipulate the host ubiquitination system through the delivery of secreted effectors known as novel E3 ubiquitin ligases (NELs). Despite the presence of NELs amongst these well-studied bacterial species, their unique structure has limited the tools that are available to probe their molecular mechanisms and explore their therapeutic potential. In this work, we report the identification of two high affinity engineered ubiquitin variants that can modulate the activity of the *Salmonella enterica* serovar Typhimurium encoded NEL, SspH1. We show that these ubiquitin variants suppress SspH1-mediated toxicity phenotypes in *Saccharomyces cerevisiae*. Additionally, we provide microscopic and flow cytometric evidence that SspH1-mediated toxicity is caused by interference with *S. cerevisiae* cell cycle progression that can be suppressed in the presence of ubiquitin variants. *In vitro* ubiquitination assays revealed that these ubiquitin variants increased the amount of SspH1-mediated ubiquitin chain formation. Interestingly, despite the increase in ubiquitin chains, we observe a relative decrease in the formation of SspH1-mediated K48-linked ubiquitin chains on its substrate, PKN1. Taken together our findings suggest that SspH1 toxicity in *S. cerevisiae* occurs through cell cycle interference and that an engineered ubiquitin variant approach can be used to identify modulators of bacterially encoded ubiquitin ligases.

**Author Summary:** Novel E3 ligases (NELs) are a family of secreted effectors found in various pathogenic gram-negative bacteria. During infection these effectors hijack vital host ubiquitin signaling pathways to aid bacterial invasion and persistence. Despite interacting with a protein as highly conserved as ubiquitin, they have a distinct architecture relative to the eukaryotic E3 enzymes. This unique architecture combined with the indispensable role ubiquitin signaling plays in host cell survival has made hindering the contribution of NELs to bacterial infections a difficult task. Here, we applied protein engineering technology to identify two ubiquitin variants (Ubvs) with high affinity for SspH1, a *Salmonella*-encoded NEL. We provide evidence that these high affinity Ubvs suppress a known SspH1-meidated toxicity phenotype in the eukaryotic model system *Saccharomyces cerevisiae*. We also show that this suppression occurs without interfering with host ubiquitin signaling. Furthermore, we demonstrate the ability of a Ubv to modulate the activity of SspH1 *in vitro*, ultimately altering the lysine linkages found in SspH1-mediated ubiquitination. To our knowledge, this is the first evidence that an engineered ubiquitin variant approach can be implemented to modulate the activity of a family of previously untargetable bacterial-encoded E3 ligases.

## Introduction

*Salmonella enterica* serovar Typhimurium is a facultatively anaerobic, rod-shaped, Gram-negative bacteria known to cause self-limiting gastroenteritis (1). It is estimated that 150 million cases of gastroenteritis linked to non-typhoidal *Salmonella* (NTS) infection occur annually worldwide, leading to an estimated 60 000 deaths (2). *S.* Typhimurium commonly invades the human host through the gastrointestinal tract, where it must cross the intestinal epithelium to establish an infection (3). Successful invasion requires *Salmonella* to induce its uptake into non-phagocytic cells, disrupt the host immune response, and assemble a *Salmonella*-containing vacuole (SCV) to act as a replication niche (4,5). *Salmonella* accomplishes these vital process using effectors which are secreted into the host cell by two type III secretion systems (T3SS) which are each encoded on a *Salmonella* pathogenicity island (SPI) (6).

A unique family of effectors secreted by these T3SS are the so-called Novel E3 Ligases (NELs) that modulate host ubiquitination to interfere with host cell signaling (7,8). These effectors are found in a multitude of pathogenic bacteria such as *Salmonella enterica, Shigella flexneri, Sinorhizobium fredii, Ralstonia solanacearum* that target a variety of eukaryotic hosts (8,9).

NELs represent a unique architecture of E3 ubiquitin (Ub) ligase, as it does not share any sequence or structural similarity to the previously described eukaryotic E3 ligases, although they are known to form a thioester bond, through a catalytic cysteine residue, to the C-terminal diglycine motif of ubiquitin in a mechanism similar to HECT eukaryotic E3 ligases (10).

Interestingly, they have evolved this architecture in the context of a prokaryotic cell, where there is an absence of ubiquitin encoding genes, indicating their function is uniquely suited to alter the host ubiquitome (11). NELs have two major domains; the N-terminal leucine rich repeat (LRR) domain and the eponymous C-terminal NEL domain. The latter harbors a catalytic cysteine and mediates the interaction with the incoming E2∼Ub conjugate, while the LRR domain mediates substrate recognition and plays a role in controlling NEL activity by preventing access to the catalytic cysteine through the adoption of an autoinhibitory conformation (7,12,13). Given that NELs play a relevant role in bacterial pathogenesis, but are structurally and mechanistically distinct from their mammalian counterparts, they represent putative pharmacological targets (8). Inhibition of these enzymes would have the advantage of not limiting the growth of the bacteria outside of the context of host infection, lowering the pressure for resistance to arise.

*Salmonella* secreted protein H1 (SspH1) was the first NEL identified in *Salmonella* and is one of four NELs encoded by *S.* Typhimurium (8,14). SspH1 is secreted by both the T3SS-1 and T3SS- 2 during infection of intestinal epithelial cells, as well as macrophages. Upon entering a host cell, it localizes to the nucleus through unknown mechanisms (14–17). Functionally, SspH1 downregulates NF-κB activity, reduces IL-8, IL-6 and CCL5 pro-inflammatory cytokine secretion, and ubiquitinates the serine/threonine protein kinase N1 (PKN1) leading to its degradation (13,15,16,18). Previous research has indicated that SspH1-mediated degradation of PKN1 can interfere with its role in potentiating the androgen receptor (AR) but is insufficient to alter AKT signaling during *Salmonella* infection. Additionally, the presence of SspH1 contributes to *Salmonella* survival during inflammatory conditions by influencing chemotaxis during infection (13,19–21).

In this study we have used a phage-displayed ubiquitin variant (Ubv) library to isolate high-affinity binders of SspH1. This technique has previously yielded modulators of human E1, E2, and E3 enzymes, as well as both human, and viral deubiquitinating enzymes (22–25). We identified two high-affinity binders, Ubv A06 and Ubv D09, which attenuated SspH1-mediated toxicity in *Saccharomyces cerevisiae* (13). Microscopic and cytometric analyses of *S. cerevisiae* growth in the presence of SspH1 revealed severe cell cycle perturbations that were relieved when these Ubv were present. *In vitro* ubiquitination assays confirmed that Ubv D09 modulated SspH1 E3 ubiquitin ligase activity, by unexpectedly increasing overall E3 ubiquitin ligase activity, while decreasing specific ubiquitin linkage types. Taken together, we provide evidence that an Ubv approach can generate effective modulators of NELs both *in vivo* and *in vitro*.

## Methods

### Cloning & Transformation

Strains and plasmids used in this study are listed in Table 1.

**Table 1.** Strains Used for Cloning.

| Name | Strain # | Description or Reference |
| --- | --- | --- |
| **Plasmids** |  |  |
| pcDNA3::2xHA-SspH1 | AB 63 | (59) |
| p426GALL::2xHA-SspH1 | AB 175 | This Work |
| pcDNA3::2xHA-SspH1<sup>C492A</sup> | AB 240 | This Work |
| p426GALL::2xHA-SspH1<sup>C492A</sup> | AB 242 | This Work |
| pDONR::Ubv A06 | APB 1 | (32,33) |
| pDONR::Ubv D09 | APB 2 | (32,33) |
| pDONR::ΔDiGly | APB 3 | (32,33) |
| pGREG515 | APB 4 | (26) |
| pDEST527 | APB 293 | pDest-527 was a gift from Dominic Esposito (Addgene plasmid #11518) |
| pDEST527::Ubv D09 | APB 300 | This Work |
| pDEST527::ΔDiGly | APB 302 | This Work |
| pGEX-PP-3xHA::SspH1 | AB 287 | This Work |
| *Escherichia coli* |  | BL21 (DE3) NEB (Cat # C2527) |
| *Saccharomyces cerevisiae* | | Wild type BY4742 $\alpha$ yeast |
| pGREG515::Ubv A06 | APB 52 | This Work |
| pGREG515::Ubv D09 | APB 53 | This Work |
| pAG426GALL::SspH1<br>+ pGREG515::ΔDiGly | APB 173 | This Work |
| pAG426GALL::SspH1<br>+ pGREG515::Ubv A06 | APB 174 | This Work |
| pAG426GALL::SspH1<br>+ pGREG515::Ubv D09 | APB 175 | This Work |
| pAG426GALL::SspH1<sup>C429A</sup><br>+ pGREG515::ΔDiGly | APB 176 | This Work |
| pAG426GALL::SspH1<br>+ pGREG515 | APB 185 | This Work |

#### Yeast Transformation

*Saccharomyces cerevisiae* (BY4742 α) were grown overnight at 30°C with shaking in complete supplement mixture (CSM) liquid media [6.7 g/L complete supplement media (with appropriate auxotrophic selection), 50 g/L ammonium sulfate, 17 g/L yeast nitrogen and 1% Glucose. Overnight cultures were harvested, washed with sterile mqH2O, washed with100 mM LiAc and resuspended in 50% (w/v) PEG 3500, 1 M LiAc, Salmon Sperm DNA (SSDNA), and 20 ng/µL of plasmid DNA. Transformations were incubated at 30°C for 30 minutes, heat shocked at 42°C for 20 minutes, harvested and resuspended in sterile mqH2O. Transformed cells were then plated on CSM plates lacking the appropriate amino acids for auxotrophic selection and incubated at 30°C.

#### Ubv Drag & Drop Cloning

Yeast expression clones of ΔDiGly, Ubv A06 and Ubv D09 were generated as described (26). In short, pGREG515 was digested with *Sal*I to expose the rec1 and rec2 sites. Rec1 and rec2 overhangs were added to ΔDiGly, Ubv A06 and Ubv D09 by PCR from the corresponding pDONR templates using primers: pGREG515UbvFor (5’- gcgtgacataactaattacatgactcgaggtcgacccaactttgtacaagaaagctggg-3’) and pGREG515UbvRev (5’- gcgtgacataactaattacatgactcgaggtcgacccaactttgtacaagaaagctggg-3’). Homologous recombination of the PCR fragment into the digested pGREG515 backbone occurred through co-transformation of *S. cerevisiae*.

#### Generation of SspH1^C492A^

The active site mutation (C492A) of SspH1 was generated in the pcDNA3::2xHA-SspH1 backbone using the Quikchange II site-directed mutagenesis kit according to the manufacturer’s protocol (Agilent). Briefly, the pcDNA3::2xHA-SspH1 template was amplified using mutagenic primers: SspH1C492AFor (5’-gcaacagaggcaacatcaactgcagagg accgggtcacacatgc-3’) and SspH1C492Arev (5’-gcatgtgtgacccggtcctctgcagttgatgttgcctctgttgc-3’) and cycling conditions suggested by the manufacturer. Amplified products were *Dpn*I-digested, transformed into DH10B *E.coli* using standard methods and sequence verified.

#### SspH1 Restriction Cloning

Yeast expression clones of SspH1 and SspH1^C492A^ were generated by introducing *Hind*III and *Xho*I fragments from pcDNA3::2xHA-SspH1 or pcDNA::2xHA- SspH1^C492A^ into p426GALL digested with the same enzymes.

#### Protein Purification cloning

Bacterial protein expression clones were generated using the Gateway® recombinational cloning system. (Invitrogen, ThermoFisher Scientific) (27,28). Briefly, pDONR::Ubv D09 and pDONR::ΔDiGly served as entry clones, which were recombined into pDEST 527 (Addgene; Plasmid #11518, Kindly donated by Dominic Esposito) according to the manufacturer’s specifications. Gateway reactions were transformed into DH5α *E. coli* before transforming BL21(DE3) *E. coli* for protein expression.

### Saccharomyces cerevisiae growth

Co-transformed yeast were grown overnight with shaking at 30°C in CSM-LEU-URA + 1% Glucose. Cells were washed 3x with sterile mqH_2_0 and resuspended in CSM-LEU-URA supplemented with either 1% Glucose (Non-inducing condition) or 1% Galactose (Inducing condition) and diluted to an OD_600_ of 1. For growth on solid medium, a 1:10 dilution series was spotted on plates containing CSM-LEU-URA + 1% Glucose or CSM-LEU-URA + 1% Galactose solid media and incubated at 30°C for 48 hours. Yeast were enumerated at the lowest concentration where growth was seen and a toxicity index was generated using the following equation: 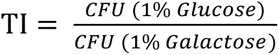 . For growth in liquid media, yeast were diluted to a starting OD_600_ of 0.1 and grown in triplicate in a 96-well plate at 30°C. The OD_600_ was measured every 10 minutes over a period of 48 hours using a Spectramax i3x Microplate Reader. Relative growth was calculated using the following equation: 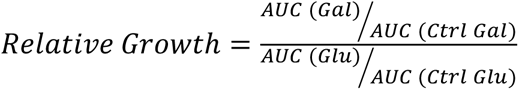 refers to either ΔDiGly or SspH1^C492A^ + ΔDiGly for figures 2 and 3, respectively.

### Flow Cytometry

Co-transformed yeast were grown overnight with shaking at 30°C in CSM-LEU-URA + 1% Glucose. 6 x 10^6^ cells/mL were harvested, washed with sterile mqH20 and resuspended in CSM-LEU-URA + 1% Galactose with 15 µg/mL of nocodazole to induce G2/M cell cycle arrest.

Cultures were grown with shaking at 30°C for 3 hours, washed with sterile mqH_2_0 and resuspended in CSM-LEU-URA + 1% Galactose. A time zero sample was removed and the remaining cultures were grown with shaking at 30°C for 480 min. Samples were harvested and resuspended in cold 70% EtOH and stored at 4°C. A 3X volume of 50 mM sodium citrate was added, cells were harvested and resuspended in 50 mM sodium citrate, 0.1 mg/mL RNase A. Cells were incubated at 37°C for 2 hours and propidium iodide in sodium citrate was added to a final concentration of 4 µg/mL. 100 000 cell events were recorded by Attune NxT and data was analyzed with FlowJo V10.6.0. Relative amount of 2N yeast was calculated using the following equation: 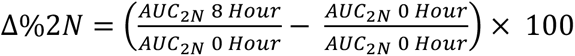

#### Microscopy

Co-transformed yeast were grown overnight with shaking at 30°C in CSM-LEU-URA + 1% Glucose. Cells were harvested, resuspended in CSM-LEU-URA supplemented with either 1% Glucose (Non-inducing) or 1% Galactose (Inducing) and grown for 8 hours with shaking at 30°C. Samples were fixed with 4% Paraformaldehyde (PFA) for 30-45 minutes at room temperature, harvested and washed 3x in PBS. Pellets were resuspended in 0.2% Triton X-100, and incubated at 4°C overnight in the dark. Samples were treated with 5.7 µM DAPI and incubated another hour in the dark at room temperature. Stained cells were harvested, washed 3x in PBS and resuspended in Vectashield. Cell suspensions were spotted onto a glass slide, covered with a round #1.5 glass coverslip, sealed with nail polish, dried in the dark and imaged using an EVOS FL Auto at 100x magnification. The 100x oil objective lens had a numerical aperture of 1.28. Micrographs were collected and analysis of yeast budding was performed as previously described using FIJI v.2.3.0 (60–62). Large-budded cells were defined as having a bud length equal to, or greater than, 1/3 of the mother cell. Analysis was performed by a person blinded to the protein expression plasmids but familiar with fluorescent microscopy acquisition methods.

### *In silico* Protein-Protein Interaction Predictions

Predicted protein structures were generated via Alphafold multimer. Molecular graphics and analyses performed with UCSF Chimera V1.14, developed by the Resource for Biocomputing, Visualization, and Informatics at the University of California, San Francisco, with support from NIH P41-GM103311 (29).

### Protein Purification

BL21 DE3 *E. coli* containing either the pDEST527 + Ubv ΔDiGly or pDEST527 + Ubv D09 were grown overnight with shaking at 37°C in Luria-Bertani broth (LB; 10 g/L tryptone, 5 g/L yeast extract,10 g/L NaCl) supplemented with 0.1 mg/mL ampicillin (Amp). Overnight cultures were subcultured 1:10 in fresh LB-AMP and incubated for 1 hour at 37°C. 400 µM isopropyl ß- D-1-thiogalactopyranoside (IPTG) was added and cells were incubated 4 hours at 37°C with shaking. Cells were harvested, resuspended in cold lysis buffer [200 mM NaPO_4_ pH 7.4, 500 mM NaCl, 25 mM imidazole, 10 µg/mL DNase A, 1 µg/mL RNase, 1x Pierce Protease Inhibitor cocktail (ThermoFisher Scientific)] prior to lysis by three passages through a French pressure cell at 1100 PSI. Lysate was successively centrifuged for 15 min at 4°C at 8000 and 30 000 x *g* then passed through a 0.45 µm filter. Nickel-NTA affinity chromatography was performed using an AKTA GO and HisTrapFF 1ml columns (Cytiva Life Sciences). Elution was performed using a 25 mM - 500 mM gradient of imidazole over 20 column volumes that also contained 20mM NaPO_4_ pH 7.4, 500mM NaCl. 0.5 mL fractions were collected, analyzed by SDS-PAGE, pooled and further purified by size exclusion chromatography (Superdex 200 Increase 10/300 GL, Cytiva Life Sciences). Fractions were collected using isocratic elution with 50 mM Tris pH 8.0, 100 mM NaCl, 1mM EDTA, 1mM DTT over 2 CV. Appropriate fractions were pooled after SDS-PAGE analysis, concentrated using a 5 kDa molecular wight cut off (MWCO) concentrator (Amicon Ultra) and refined by removing higher molecular weight species via a 30 kDa MWCO concentrator (Amicon Ultra).

Recombinant SspH1 with tandem N-terminal GST and HA epitope tags was purified according to the procedure outlined in (30) before PreScission protease digestion to remove the GST tag. Briefly, 50 µg of purified GST-HA-SspH1 was mixed with 2 µg in-house purified GST-tagged PreScission protease and pre-equilibrated Glutathione Sepharose 4B beads (GE Healthcare) in PreScission protease buffer (50 mM Tris pH 7.5, 300 mM NaCl, 1 mM EDTA, 1 mM DTT). Reactions were incubated overnight to allow for protease cleavage between the GST and HA tags of SspH1 before beads were removed by centrifugation. Cleaved HA-SspH1 was recovered from the supernatant.

### Mass Spectrometry

Mass spectrometry work was performed by the Alberta Proteomics and Mass Spectrometry Facility in the Faculty of Medicine and Dentistry at the University of Alberta. Purified ubiquitin variant protein was separated on 4-20% polyacrylamide gradient gels (Bio-Rad) by electrophoresis. The gel was washed 3x with mqH_2_O, stained with Imperial protein stain (ThermoFisher Scientific) for 2 hours at room temperature with shaking and destained overnight with mqH_2_0 at room temperature with shaking. Protein bands of interest were excised, reduced (10 mM β-mercaptoethanol in 100 mM ammonium bicarbonate) and alkylated (55 mM iodoacetamide in 100 mM ammonium bicarbonate). After dehydration enough trypsin (6ng/ul, Promega Sequencing grade) was added to just cover the gel pieces and the digestion was allowed to proceed overnight (∼16 hrs.) at 37°C. Tryptic peptides were first extracted from the gel using 97% H_2_O, 2% acetonitrile, 1% formic acid followed by a second extraction using 50% of the first extraction buffer and 50% acetonitrile.

The tryptic peptides were resolved using nano flow HPLC (Easy-nLC 1000, Thermo Scientific) coupled to an Orbitrap Q Exactive mass spectrometer (Thermo Scientific) with an EASY-Spray capillary HPLC column (ES902A, 75 um x 25 cm, 100 Å, 2 µm, Thermo Scientific). The mass spectrometer was operated in data-dependent acquisition mode with a resolution of 35,000 and m/z range of 300–1700. The twelve most intense multiply charged ions were sequentially fragmented by using HCD dissociation, and spectra of their fragments were recorded in the orbitrap at a resolution of 17,500. After fragmentation all precursors selected for dissociation were dynamically excluded for 30 s. Data was processed using Proteome Discoverer 1.4 (Thermo Scientific) and the database was searched using SEQUEST (Thermo Scientific). Search parameters included a strict false discovery rate (FDR) of .01, a relaxed FDR of .05, a precursor mass tolerance of 10 ppm and a fragment mass tolerance of 0.01 Da. Peptides were searched with carbamidomethyl cysteine as a static modification and oxidized methionine and deamidated glutamine and asparagine as dynamic modifications.

### SspH1 Ubiquitination Assays

0.15 µg of purified HA-SspH1 was incubated with 0.22 µg of recombinant human UBE1, 4.0 µg of human UBE2D2, 1.8 µg of HA-ubiquitin (all R&D Systems), His-ΔDiGly, His-Ubv D09, and/or 0.41 µg GST-PKN1 (ThermoFischer Scientific) in ubiquitination reaction buffer (80 mM Tris-HCl pH 7.6, 50 mM NaCl, 10 mM MgCl2, 0.1 mM DTT) and initiated with 2 mM ATP. All samples were incubated for 3 hours at 37°C prior to being quenched by addition of SDS-PAGE sample buffer and boiling at 100°C for 5 minutes.

### Immunoprecipitation

Ubiquitination reactions containing recombinant GST-PKN1 were immunoprecipitated using protein G-conjugated magnetic beads (New England Biolabs). Beads were prepared by washing three times with IP wash buffer (PBS + 0.1% Tween-20), then incubated with 2 µg Rabbit α-GST polyclonal antibody (Santa Cruz Biotechnology; Cat #sc-459) for 20 minutes at room temperature with agitation. Washing steps were repeated to remove unbound antibody, then beads were blocked with 3% milk powder solution for 1 hour at 4°C. After blocking, beads were washed again, and ubiquitination reaction samples were incubated with the beads for 1 hour at 4°C with agitation. Washing steps were performed a final time followed by elution from the beads by addition of SDS-PAGE sample buffer and boiling at 100°C for 5 minutes.

### Immunoblotting

Proteins were separated by polyacrylamide gel electrophoresis and transferred to nitrocellulose membranes (Bio-Rad). Membranes were dried, rehydrated with Tris-buffered saline (TBS) and blocked with TBS blocking buffer (Li-Cor) before incubation with primary antibody diluted in TBS blocking buffer overnight. Membranes were washed and incubated with secondary antibodies diluted in the same buffer for 1 hour. The antibodies used in this study are: mouse α- Actin (sc-8433; Santa Cruz Biotechnology) 1: 2000; mouse α-Myc (9E10; Provided by Dr. Rob Ingham, University of Alberta) 1:2 500; mouse α-His (27E8; Cell Signaling Technology) 1: 2 500; rabbit α-K48-linkage specific polyubiquitin (D9D5; Cell Signaling Technology) 1:2 000; rabbit α-K63-linkage specific polyubiquitin (D7A11; Cell Signaling Technology) 1:2 000; mouse α-Ubiquitin (P4D1; Cell Signaling Technology) 1:2 000; rat α-HA (3F10; Roche Diagnostics) 1:2 500; rabbit α-GST (sc-459; Santa Cruz Biotechnology) 1:2 500; goat α-mouse (926-68020; Licor) 1:5 000; goat α-rabbit (925-32211; Licor) 1:5 000; goat α-rat (926-32219; Licor) 1:5 000. Blots were imaged with a Li-Cor Odyssey and visualized/band intensity quantified using Imagestudio V5.2.5.

### Statistical Analysis

All statistical comparisons were performed using Graphpad Prims 9.5.1. Data are presented as the mean with error bars representing SEM. Growth Reduction Co-efficient (GRC) for comparison of yeast growth in liquid media was calculated as described in Lauman and Dennis (31). Statistical analyses were determined through one-way ANOVA with Tukey’s multiple comparison test or unpaired t-test. Statistical significance is indicated as follows: *P*>0.05 = ns, *P*<0.05 = *, *P* <0.01 = **, *P* <0.001 = ***, *P* <0.0001 = ****.

## Results

### Ubiquitin Variants as Inhibitors of SspH1

A previously described ubiquitin variant library was used to screen for binding interactions with SspH1 (32–34). Two ubiquitin variants (Ubvs), designated A06 and D09 were identified with enhanced binding to SspH1 relative to wildtype ubiquitin (Fig. 1A). Sequence alignment revealed 12 amino acid differences between human ubiquitin and either ubiquitin variant, but only 2 amino acid differences between Ubv A06 and Ubv D09 (Fig. 1B, C). We used homology modeling to predict Ubv A06 and D09 protein structures, and UCSF Chimera to compare the spatial positioning of the altered amino acids between human ubiquitin and the ubiquitin variants (29,35). Mutations in Ubv A06 and D09 were mainly found in the Isoleucine 44 recognition patch of ubiquitin, which is a known interface in E2-E3 ubiquitin transfer, as well as in the C-terminal tail (36). Predictive and comparative structural modeling done with Alphafold Multimer revealed two predicted wild type ubiquitin binding sites on SspH1, one within the active site and a second along the C-terminal thumb domain, which is known to be the E2 interacting motif (37) (Fig 1D). Ubv A06 and D09 were also predicted to bind within these pockets suggesting the mutations do not vastly change the structural relationship between the Ubv and SspH1, which is notable since the Ile 44 patch is predicted to be the primary interaction face between the Ubv and SspH1. Collectively, these results suggest that Ubv A06 and Ubv D09 may have improved binding to SspH1 compared to human ubiquitin.

**Fig 1.**
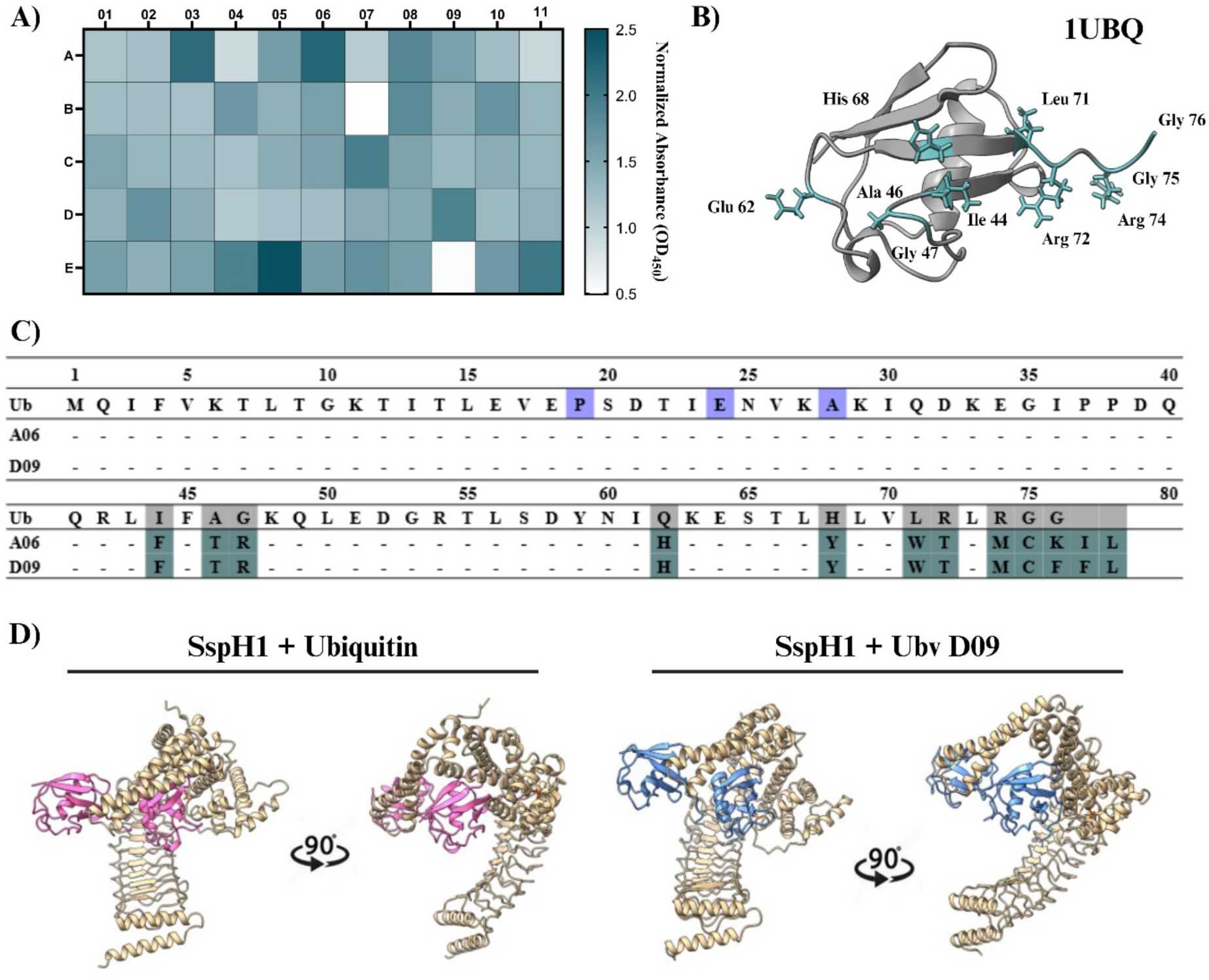
Identifying Ubiquitin Variants with a High Binding Affinity for SspH1. **(A**) The binding specificities of phage displayed Ubvs as assessed by phage ELISA. Subsaturating concentrations of phage were added to immobilized proteins as indicated. Bound phages were detected by the addition of anti-M13-HRP and colorimetric development of TMB peroxidase substrate. The mean value of the absorbance at 450 nm is indicated by color. Variant labels were based on the letter and number indicated along the y- and x-axis, respectively. **(B)** Structural depiction of human ubiquitin (1UBQ) with the mutated residues highlighted and the wildtype side chains shown. **(C)** Sequences of Ubvs that bind with a high affinity to SspH1. Amino acids differences between human ubiquitin, ubiquitin variant A06 and ubiquitin variant D09 are highlighted in green. Amino acid differences between human ubiquitin and *S. cerevisiae* ubiquitin are highlighted in purple. **(D)** Alphafold multimer predictions of SspH1 interacting with human ubiquitin, in pink, or ubiquitin variant D09, in blue. The catalytic residue of SspH1, Cys 492, is highlighted in orange. The thumb region is located at C-terminus of NEL domain.

### Ubvs A06 & D09 Suppress SspH1-Mediated Toxicity in Yeast

Yeast are a robust eukaryotic model, have well-developed genetic tools and contain all the necessary components of the ubiquitin system (38,39). We took advantage of the inducible GAL expression system to heterologously express SspH1, its catalytic variant (SspH1^C492A^) and 3 ubiquitin variants – Ubv A06, D09 and a WT construct lacking the final diglycine motif at the C-terminus (ΔDiGly). The ΔDiGly construct was used to ensure that any changes to SspH1 function were due to the difference in Ubv affinity for SspH1 and not because of their inability to form the initial thioester linkage between SspH1 and ubiquitin. An empty vector (Ev) plasmid was also expressed alongside SspH1 to determine the effect of SspH1 in the absence of Ubv. We chose to study the functional interaction of SspH1 and Ubvs in a yeast model system because it has been previously shown that catalytically active SspH1 is toxic to yeast making this a robust selection model, rather than a screen (13). Yeast ubiquitin differs from human ubiquitin at three locations, Ser19, Asp24 and Ser28, none of which are mutated in either Ubv (40) (Fig. 1B).

We confirmed that all proteins were expressed under our assay conditions (Fig. 2A). To first determine if expression of either Ubv A06 or D09 was detrimental to yeast growth, we expressed each Ubv individually and monitored yeast growth in both liquid and solid media over a 48 hour period. Growth in liquid media was quantified using the relative growth equation described by Lauman and Dennis (31). We observed similar growth of yeast expressing Ubv A06 and Ubv D09 compared to the non-inducing condition in both solid and liquid media, indicating that Ubv expression alone does not confer toxicity by interfering with the endogenous Ubiquitin-proteasome-system (Fig. 2B-E).

**Fig 2.**
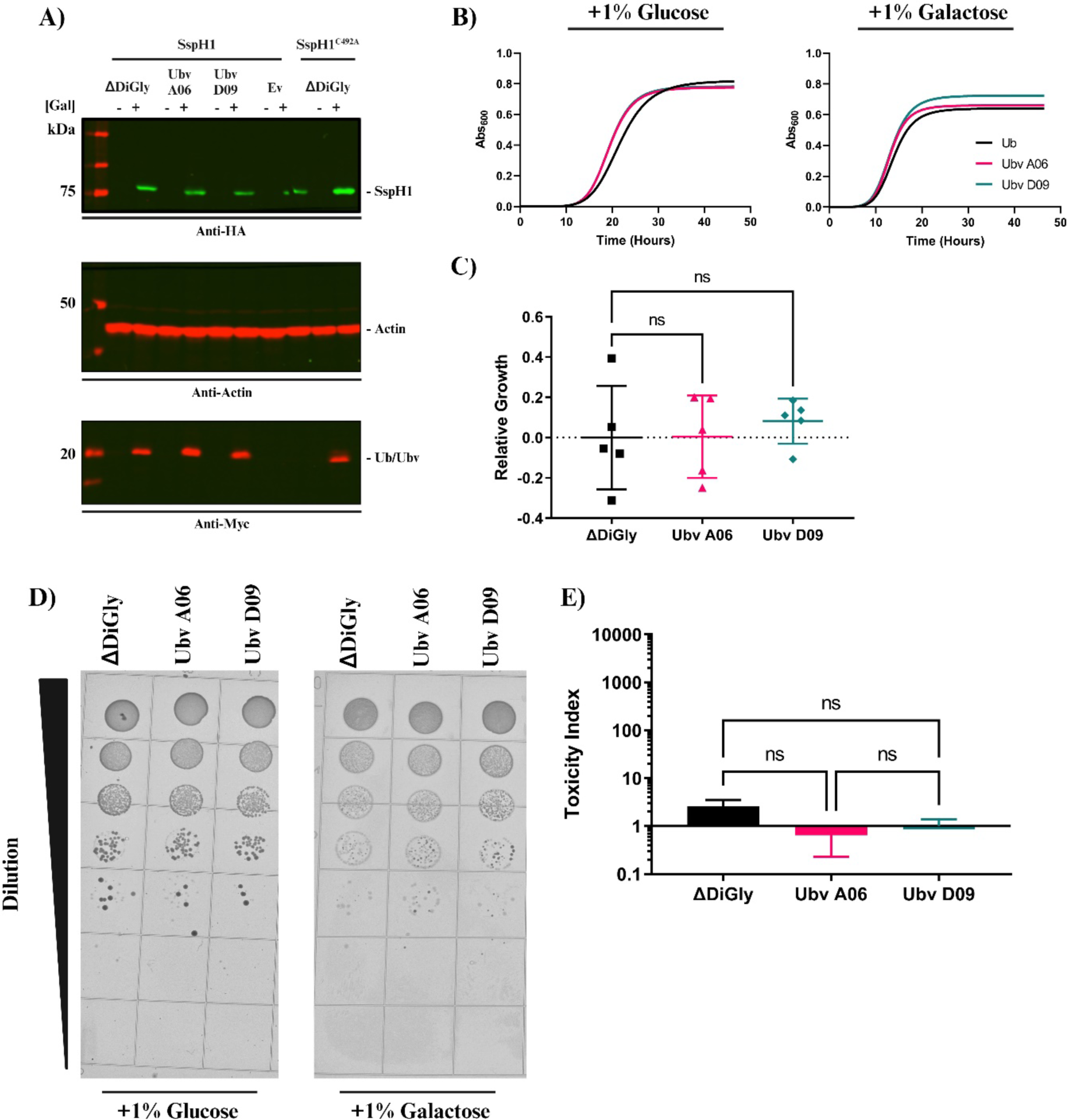
Ubv A06 & D09 are not Toxic to Yeast when Expressed Alone. **(A)** Expression of SspH1 or SspH1^C492A^ and Ubv ΔDiGly, Ubv A06 or Ubv D09 in BY4742α yeast strain co-transformed with galactose-inducible vectors (pGREG515). SspH1 was detected through the use of anti-HA staining whereas Ubvs were detected through anti-Myc staining **(B)** Growth of BY4742α yeast strain transformed with galactose-inducible Ubv ΔDiGly, Ubv A06 or Ubv D09. Strains were grown overnight in 1% glucose then washed and diluted in 1% galactose or 1% glucose, as indicated, for 48 hours at 30°C in a 96-well plate. Growth was monitored by measuring the Abs_600_ every 10 mins for the duration of the 48-hour growth period. **(C)** Quantification of strain growth using relative growth, where the area under the curve (AUC) for each strain was calculated and compared to the control (Ubv ΔDiGly) in both the inducing and non-inducing conditions. Errors bars represent the standard error of the mean across 5 independent replicates. Relative growth calculated as described in methods. Data was analyzed by one-way ANOVA using Tukey’s multiple comparisons test. **(D)** Viability of BY4742α yeast strain transformed with galactose-inducible Ubv ΔDiGly, Ubv A06 or Ubv D09. Strains were spotted as a serial dilution series on 1% galactose or 1% glucose, as indicated, and imaged after 48 hours. **(E)** Quantification of survival on solid media by toxicity index. Errors bars represent the standard error of the mean across 3 independent replicates. Toxicity Index calculated as described in methods. Data was analyzed by one-way ANOVA using Tukey’s multiple comparisons test.

Having determined that the ubiquitin variants have no detrimental effect on yeast growth when expressed alone, we next sought to determine if Ubv co-expression would have any functional consequences on SspH1 by monitoring co-expression of Ubv A06 or Ubv D09 and SspH1 in yeast (12). The baseline for yeast growth was determined in the presence of SspH1^C492A^ + ΔDiGly because SspH1 toxicity in yeast requires its E3 ubiquitin ligase activity (12,41,42) (Fig. 3A,B). Following the expression of SspH1 + Ev or SspH1 + Ubv ΔDiGly in liquid media we observed a significant decrease, ∼20% and ∼40% respectively, in the relative growth of yeast (Fig. 3A,B), which is consistent with the previously reported effect of SspH1 expression in yeast (12). By contrast, co-expression of SspH1 with Ubv A06 or Ubv D09 led to no significant difference in relative growth when compared to yeast grown in the presence of SspH1^C492A^ (Fig. 3A,B).

**Fig 3.**
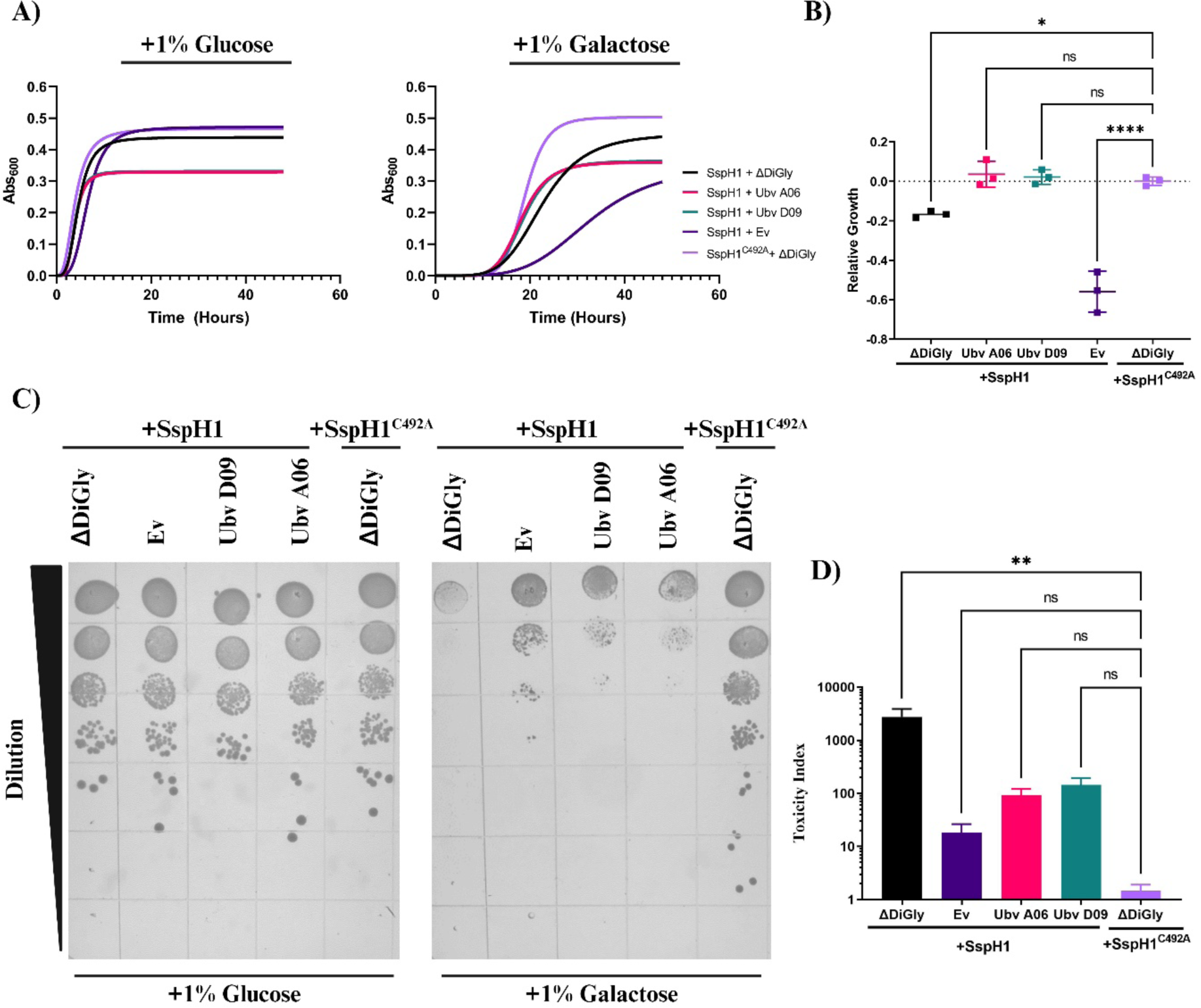
Ubv A06 & D09 Suppress SspH1-Mediated Toxicity in Yeast. **(A**) Growth of BY4742α yeast strain co-transformed with galactose-inducible SspH1 or SspH1^C492A^ and Ubv ΔDiGly, Ubv A06 or Ubv D09. Strains were grown overnight in 1% glucose then washed and diluted in 1% galactose or 1% glucose, as indicated, for 48 hours at 30°C in a 96-well plate. Growth was monitored by measuring the Abs_600_ every 10 mins for the duration of the 48-hour growth period. **(B)** Quantification of strain growth using relative growth, where the area under the curve (AUC) for each strain was calculated and compared to the control (SspH1^C492A^ + Ubv ΔDiGly) in both the inducing and non-inducing conditions. Errors bars represent the standard error of the mean across 5 independent replicates. Relative growth calculated as described in methods. Data was analyzed by one-way ANOVA using Tukey’s multiple comparisons test. **(C)** Viability of BY4742α yeast strain transformed with galactose- inducible SspH1 or SspH1^C492A^ and Ubv ΔDiGly, Ubv A06 or Ubv D09. Strains were spotted as a serial dilution series on 1% galactose or 1% glucose, as indicated, and imaged after 48 hours. **(D)** Quantification of survival on solid media by toxicity index. Errors bars represent the standard error of the mean across 3 independent replicates. Toxicity Index calculated as described in methods. Data was analyzed by one-way ANOVA using Tukey’s multiple comparisons test.

Interestingly, similar assays on solid media showed that Ubv A06 and D09 only partially suppressed SspH1 toxicity. As expected, we observed a lack of yeast toxicity in the presence of SspH1^C492A^ + ΔDiGly as well as a robust level (∼1000-fold) of toxicity in the presence of SspH1+ ΔDiGly (12) (Fig. 3C,D). This toxicity was decreased 20-fold when SspH1 was expressed alongside Ubv A06 or alongside Ubv D09 in comparison to SspH1 + ΔDiGly, although yeast growth was not rescued to baseline SspH1^C492A^ + ΔDiGly levels. Interestingly, on solid medium, co-expression of Ev with SspH1 showed robust growth rescue, unlike in liquid medium (Fig. 3C,D). Taken together, these results indicate that the presence of Ubv A06 and Ubv D09 is sufficient to suppress the SspH1-mediated toxicity of yeast growth.

### Ubv A06 & D09 Suppress SspH1-Mediated Cell Cycle Arrest in Yeast

To further elucidate the effect of ubiquitin variants on SspH1, we used flow cytometry and microscopy to examine perturbations in the yeast cell cycle caused by SspH1 expression (38,43,44). Yeast nuclei were stained with DAPI and both brightfield and fluorescent images were acquired. Direct observation of yeast co-expressing SspH1 + ΔDiGly revealed a high proportion of large-budded cells within the population that was significantly reduced in yeast co-expressing SspH1^C492A^ + ΔDiGly (Fig. 4). A similarly high proportion of large-budded cells was also observed in yeast co-expressing SspH1 + Ev (Fig. 4). This large-budded phenotype suggested that yeast toxicity may be caused by cell cycle interference leading to issues progressing through G2/M (45). Notably, the proportion of large-budded yeast was significantly reduced when SspH1 was expressed alongside Ubv A06 or Ubv D09 relative to when SspH1 was expressed alongside ΔDiGly or alone (Fig. 4).

**Fig 4.**
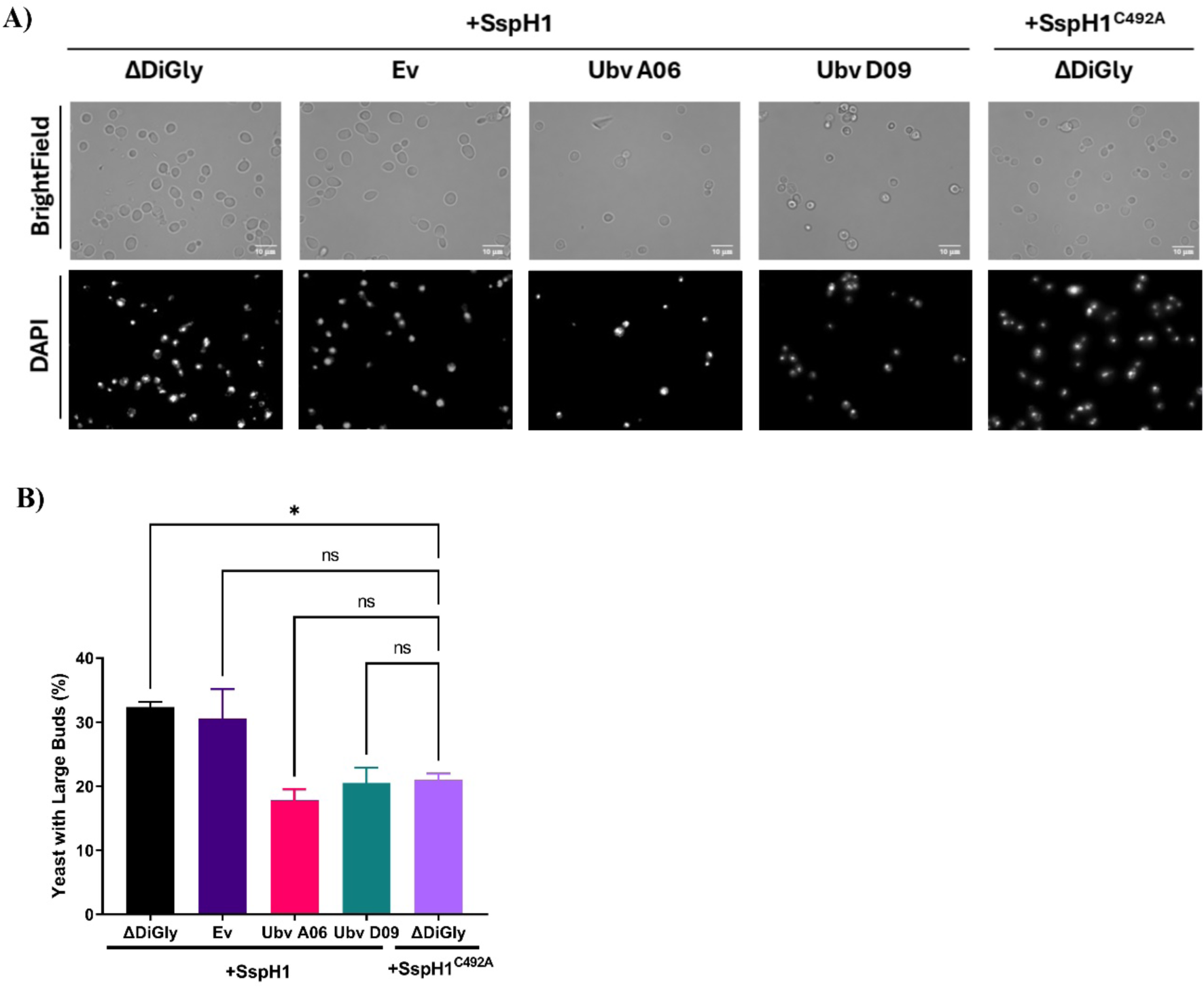
Ubv A06 & D09 Suppress SspH1-Mediated Arrest at the Large Budded Stage in Yeast. **(A)** Representative micrographs of yeast co-expressing SspH1 + ΔDiGly, Ubv A06, Ubv D09 or Ev, as well as, SspH1^C492A^ + ΔDiGly are shown after 8 hours of incubation at 30°C in fresh 1% galactose. Images were collected on an EVOS FL Auto at 100x magnification. DNA was stained in blue using 4′,6-diamidino-2-phenylindole (DAPI). **(B)** Quantification of large, budded yeast was performed as previously described using FIJI v.2.3.0 (https://fiji.sc/). (60–62) (Ex. Large budded = >1/3 mother cell size). Data was analyzed by one-way ANOVA using Dunnett’s multiple comparisons test.

As cell cycle dysregulation was implicated in SspH1-mediated toxicity in yeast, we further interrogated cell cycle dynamics through flow cytometric analyses of cellular DNA content (46). Yeast were arrested in the G2/M phase of the cell cycle using nocodazole, then released by being placed in fresh media. Escape from this arrest was measured by quantifying the proportion of 1N vs 2N DNA content (47,48). Consistent with our previous observation in the growth assays, we observed a substantial decrease (∼10-20%) in the proportion of yeast with 2N DNA content after 8 hours in yeast expressing ΔDiGly, Ubv A06, or Ubv D09 (Fig. 5A,B). This suggests that the ubiquitin variants alone are not contributing to the cell cycle interference phenotype. Similarly, in yeast expressing SspH1^C492A^ + ΔDiGly, we also observed a ∼20% decrease in yeast with 2N DNA content after 8 hour, suggesting progression through the cell cycle had resumed (Fig. 5C,D). In yeast expressing SspH1 + Ev, we only observed a ∼5% decrease in the proportion of yeast with 2N DNA content while in yeast expressing SspH1 + ΔDiGly we observed a ∼5% increase in the proportion of yeast with 2N DNA content (Fig. 5C,D). These results suggest that the presence of SspH1 prevents progression through the G2/M phase of the cell cycle. By contrast, co-expression of either Ubv allowed for progression through the cell cycle as evidenced by the ∼10-20% decrease in yeast with 4N DNA (Fig. 5C,D).

**Fig 5.**
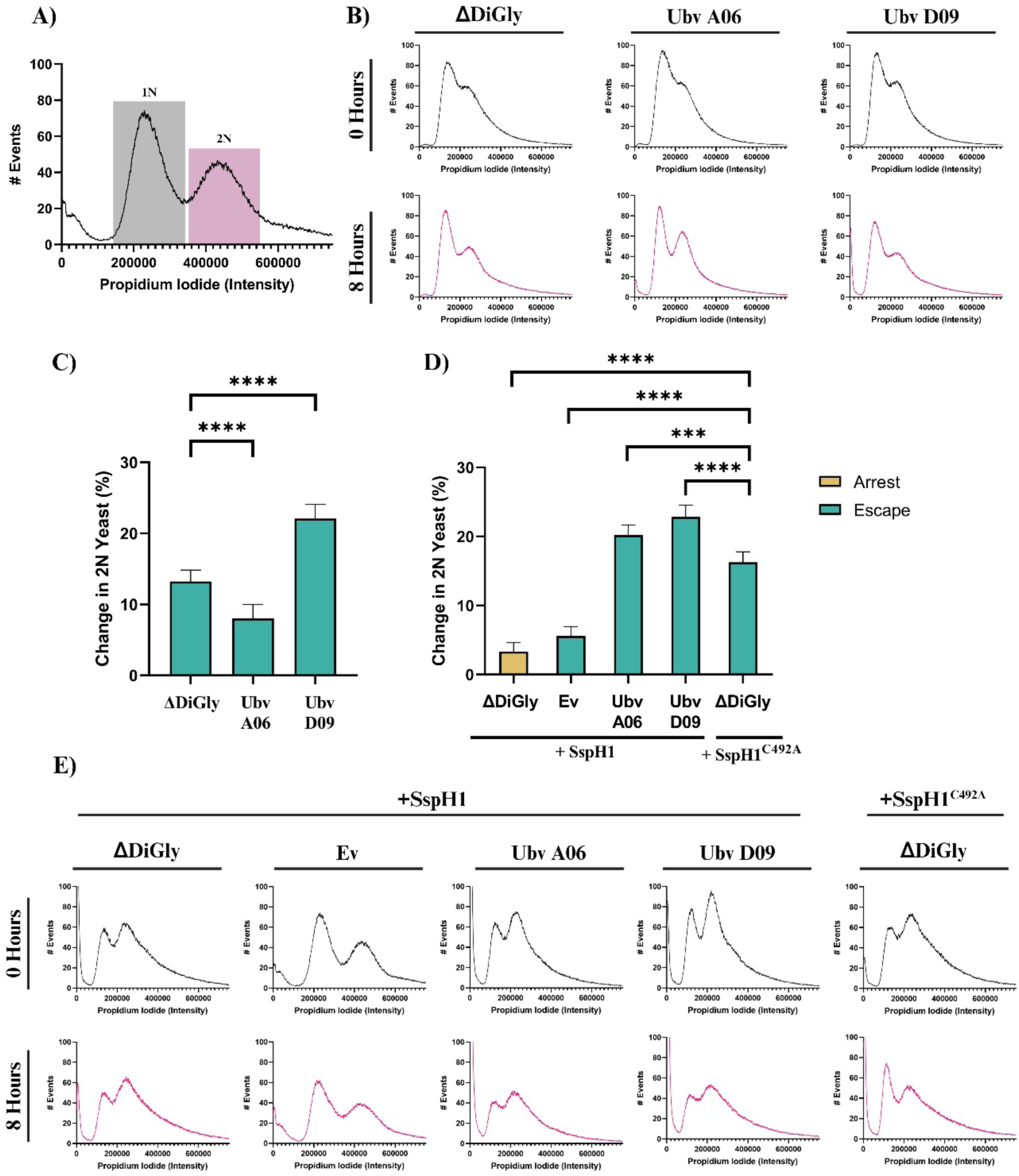
Ubv A06 & D09 Suppress SspH1-Mediated Cell Cycle Arrest in Yeast. **(A)** Example of regions used to calculate AUC for yeast with 1N (G1) and 2N (G2/M) DNA content **(B)** Cell cycle analysis of BY4742α yeast strain transformed with galactose-inducible Ubv ΔDiGly, Ubv A06, or Ubv D09. Cell cycles were synchronized at the G2/M phase through treatment with 20 *µ*M nocodazole for 3 hours at 30°C than washed multiple times to allow yeast to progress through cell cycle. Yeast were placed into fresh 1% galactose and incubated at 30°C for 8 hours prior to being fixed and having their DNA content stained with propidium iodide (PI). A 0 hour sample was also obtained immediately following the removal of nocodazole. (**C, D)** Quantification of the relative change of yeast arrested with 2N DNA content was calculated as the area under the curve (AUC) of the 2N peak at 8 hours relative to the AUC of the 2N peak immediately after nocodazole release as described in the methods. Data are shown as mean ± SEM of N=11 replicates (**C**) or N=5 replicates (**D**). Data was analyzed by one-way ANOVA using Dunnett’s multiple comparisons test. **(E)** Cell cycle analysis as described above of BY4742α yeast strain co-transformed with galactose-inducible SspH1 or SspH1^C492A^ and Ubv ΔDiGly, Ubv A06, Ubv D09 or Empty Vector (Ev).

Together these results suggest that SspH1-mediated toxicity may be caused by cell cycle interference, specifically the inability to progress through the G2/M phase of yeast budding. These results also suggest that ubiquitin variants A06 or D09 are sufficient to suppress the SspH1-mediated block in cell cycle progression.

### Ubiquitin variants alter SspH1 E3 ubiquitin ligase activity *in vitro*

As SspH1 toxicity phenotypes in yeast were not observed with a catalytic mutant, its E3 ubiquitin ligase activity is likely involved. Accordingly, we next investigated the effect of Ubv D09 on SspH1 E3 ubiquitin ligase activity. Despite multiple purification approaches, we were unable to purify Ubv A06 for use in these recombinant assays. We cannot rule out that Ubv A06 expression may be toxic to BL21 DE3 *E. coli*, although it is unclear why this would be given the sequence similarity to Ubv D09. We performed *in vitro* ubiquitination assays with recombinant purified proteins as previously described, except with the addition of purified Ubv D09 or Ubv ΔDiGly (7,12,49) (S1 Fig.). In this assay, SspH1 activity is assessed by the presence and intensity of high molecular weight ubiquitin chains in the presence of E1 and E2 enzymes, as well as ATP. As expected, we did not observe any high molecular weight His-ubiquitin chains when SspH1 was provided His-Ubv D09 as the sole ubiquitin source, since it lacks the di-glycine motif at the C-terminus (50) (Fig. 6A). Additionally, we did not observe any high molecular weight His-ubiquitin chains when SspH1 was provided both HA-ubiquitin and His-Ubv D09, indicating that Ubv D09 is not incorporated into any ubiquitin chains (Fig. 6A). However, high molecular weight HA-ubiquitinated species were readily observed, suggesting that the presence of Ubv D09 does not abrogate the ability of SspH1 to form ubiquitin chains (Fig. 6B). When SspH1 activity was assessed in the presence of ΔDiGly we observed a slight decrease in the amount of high molecular weight HA-ubiquitinated species when compared to SspH1 activity provided with only HA-ubiquitin (Fig. 6C,D). Unexpectedly, in the presence of Ubv D09, we observed a significant, ∼2.5 fold increase in the amount high molecular weight HA-ubiquitinated species relative to HA-ubiquitin alone (Fig. 6C,D). These results confirm that Ubv D09 can potentiate SspH1 E3 ubiquitin ligase activity without itself being a productive substrate.

**Fig 6.**
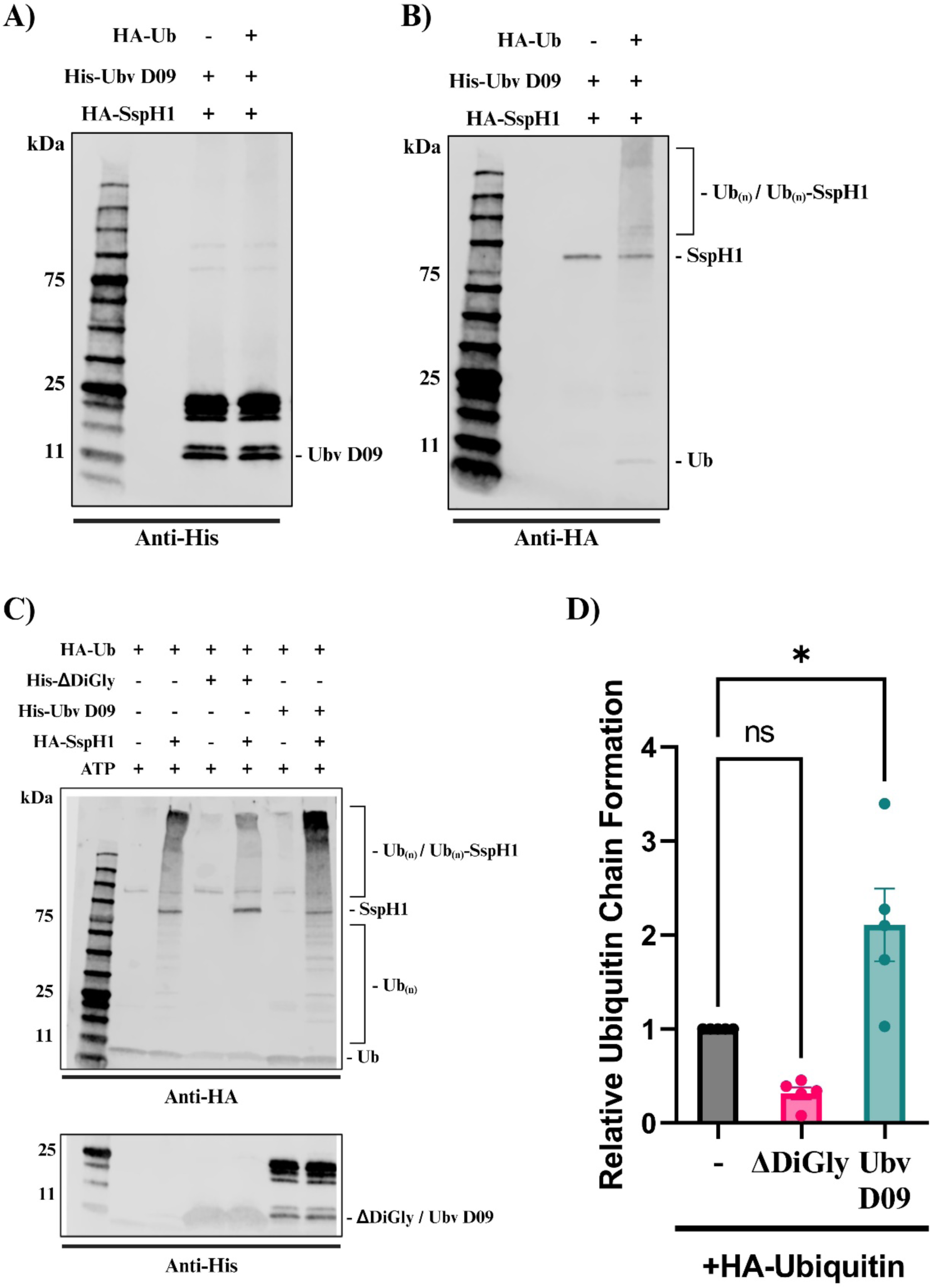
Ubvs modulate the ubiquitination activity of SspH1 *in vitro*. **(A)** The ability of Ubv D09 to be incorporated into SspH1-mediated ubiquitination was determined by *in vitro* ubiquitination assays containing recombinant E1, E2, SspH1, Ubv and ATP with or without HA-Ub as indicated (-/+). SspH1 activity was analyzed with incorporation of Ubv D09 being monitored by anti-His immunoblot. (**B**) Ubv D09 impact on polyubiquitin chain formation under the same conditions was monitored by anti-HA immunoblot. Species of interest are indicated on the right. **(C)** The effect of Ubv D09 on the ubiquitination activity of SspH1 was assessed by *in vitro* ubiquitination assays containing recombinant E1, E2, SspH1, HA-Ub, Ubv ΔDiGly, or Ubv D09 as indicated (-/+). SspH1 activity was analyzed with Ubv detected by anti-His immunoblot (Bottom) and polyubiquitin chain formation as well as SspH1 detected by anti-HA immunoblot (Top). Species of interest are indicated on the right. **(D)** HA-ubiquitin chain amount was determined through the addition of HA signal in the indicated areas of the immunoblot (Ub_(n)_ + Ub_(n)_-SspH1) and is presented as a ratio of SspH1 + HA-Ub signal. Errors bars represent the standard error of the mean across 4 independent experiments. Data was analyzed by one- way ANOVA using Tukey’s multiple comparisons test.

Given that Ubv D09 appeared to potentiate SspH1 activity, we tested if the presence of Ubv D09 alters the ubiquitination pattern of PKN1, a known SspH1 substrate (13,15). To do this, we conducted *in vitro* ubiquitination assays, as outlined above, in the presence of PKN1 and isolated PKN1 species by immunoprecipitation. As expected, in the absence of SspH1, we did not observe an upwards shift in molecular weight for PKN1, indicating a lack of PKN1 ubiquitination (Fig. 7A). In the presence of SspH1, we observed the formation of high molecular weight species which correspond to ubiquitinated PKN1, confirming that SspH1 was capable of ubiquitinating PKN1 *in vitro* (7) (Fig. 7A). We observed no significant change in the relative amount of ubiquitinated PKN1 upon addition of Ubv ΔDiGly (Fig. 7A,B). Consistent with our previous results, the addition of Ubv D09 led to a significant ∼2-fold increase in the amount of ubiquitinated PKN1 (Fig. 7A,B). Together these results suggest that Ubv D09 has a potentiating effect on the ability of SspH1 to ubiquitinate a known substrate, PKN1, *in vitro*.

**Fig 7.**
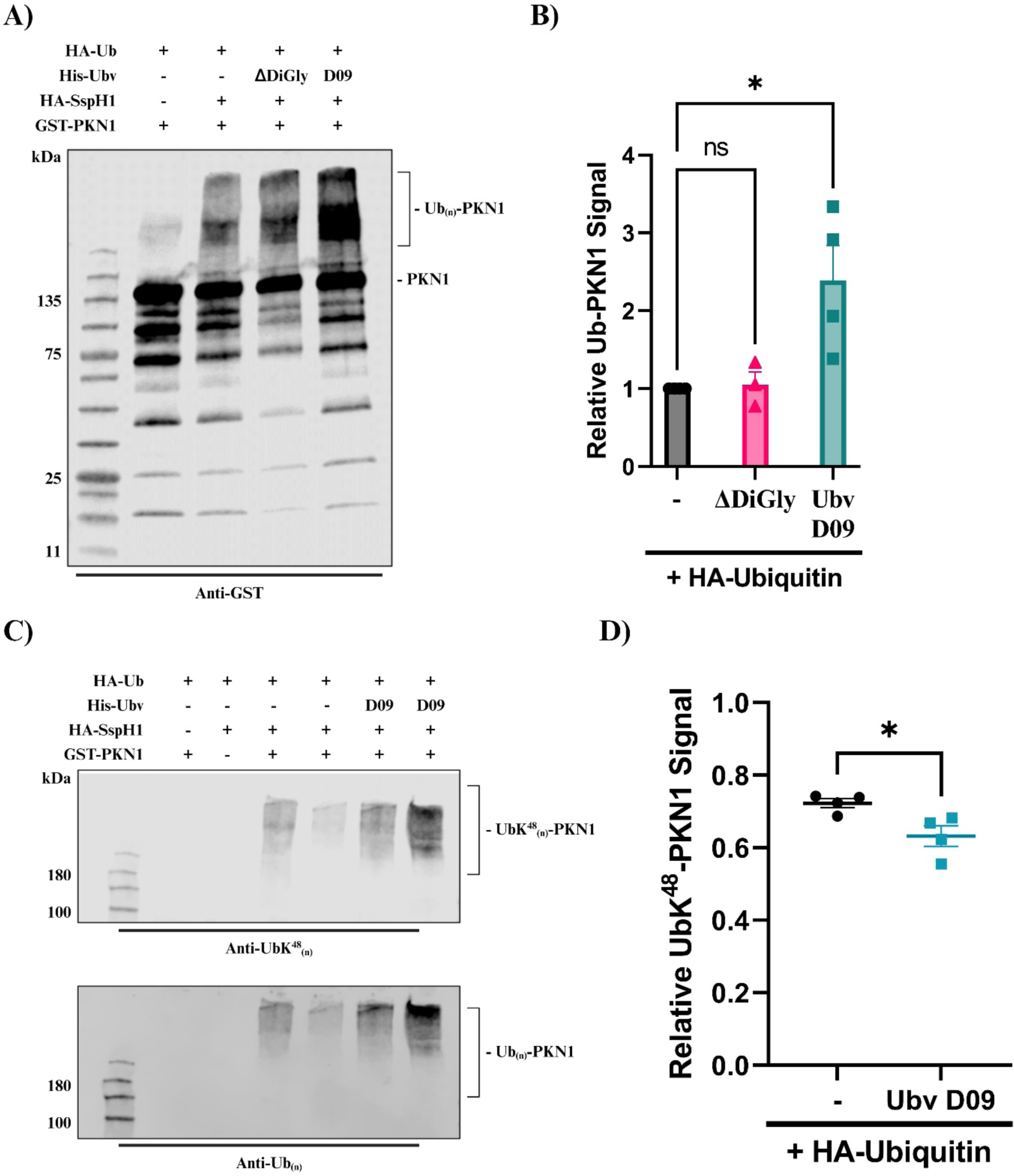
Ubvs modulate SspH1-mediated ubiquitination of PKN1 *in vitro*. **(A)** SspH1-mediated ubiquitination of PKN1 was determined by *in vitro* ubiquitination assays containing recombinant E1, E2, SspH1, PKN1, HA-Ub, Ubv ΔDiGly, or Ubv D09 as indicated. Formation of Ub_(n)_-PKN1 was monitored using anti-GST immunoblot (PKN1 has GST fusion). Species of interest are indicated on the right. **(B)** Formation of Ub_(n)_-PKN1 is expressed as ratio relative to SspH1 + HA-Ub. Error bars represent the standard error of the mean across 4 independent experiments. Data was analyzed by one-way ANOVA using Dunnett’s multiple comparisons test. **(C)** Lysine-specific ubiquitin chain conformation of PKN1-specific, SspH1- mediated ubiquitination was determined by *in vitro* ubiquitination assays containing recombinant E1, E2, SspH1, PKN1, HA-Ub, or Ubv D09 as indicated and analyzed by immunoblot. Two independent reactions are shown. Total ubiquitination was determined by anti-Ub_(n)_ [P4D1], K48-specific ubiquitin chains was determined by anti-Ub^K48^ [D9D5] **(D)** Formation of Ub^K48^ -PKN1 and Ub -PKN1 is expressed as ratio relative to the signal of Ub -PKN1. Error bars represent the standard error of the mean across 4 independent experiments. Data was analyzed using an unpaired T-test.

The suppressive effect of Ubv D09 on SspH1 toxicity in yeast led us to hypothesize that SspH1 E3 ubiquitin ligase activity was compromised, but our recombinant protein studies suggested this was not the case. To reconcile these observations, we assessed any potential differences in ubiquitin linkage which could impact substrate fate in the cell. Accordingly, we performed the previously described ubiquitination reactions followed by an immunoprecipitation to isolate PKN1 then probed with antibodies specific for Lys48- and Lys63-linked ubiquitin chains, as well as global ubiquitin antibody, to uncover the relative amount of Lys48- and Lys63-linked ubiquitin chains that were present on PKN1 (51). In the absence of SspH1 or PKN1 there was no observable Lys48, Lys63 or non-lysine specific ubiquitin chain formation (13) (Fig. 7C). When both SspH1 and PKN1 were present we observed PKN1-specific ubiquitin chain formation with ∼75% of the total ubiquitin chains being Lys-48 specific (52) (Fig. 7C,D). The addition of Ubv D09 led to an increase in the overall amount of ubiquitination we detected, which was consistent with our previous experiments (Fig. 7C,D). Interestingly, we also observed a small but significant decrease in the amount of Lys-48 specific ubiquitin chains in the presence of Ubv D09, which accounted for only ∼65% of the total ubiquitin chains, representing a 13% decrease in the relative amount of PKN1-specific Lys-48 ubiquitin chains in the presence of HA-ubiquitin alone (Fig. 7C,D). We did not observe the formation of Lys63-linked ubiquitin chains in the presence of HA-ubiquitin or HA-ubiquitin + Ubv D09 (S2 Fig.). Taken together these results suggest that, although the presence of Ubv D09 leads to an overall increase in PKN1 ubiquitination, it may interfere with the ability of SspH1 to form Lys48-linked ubiquitin chains.

## Discussion

In this study we provide evidence that modulators of bacterial ubiquitin ligase activity can be found within a ubiquitin variant library that was designed to target human ubiquitin-interacting proteins (33). We report the identification of two high-affinity Ubv binders, Ubv A06 and Ubv D09, to SspH1, one of four *Salmonella*-encoded NELs. Both Ubvs contain 12 mutations which are not conserved in human and yeast ubiquitin and the Ubvs differed from each other by only two amino acids. The mutated residues reside exclusively in diversified regions 2 and 3 of the ubiquitin variant library. *In silico* protein-protein interaction prediction suggests that the Ubvs interact with both the active site and E2∼Ub binding site of SspH1. This is consistent with the observed binding interactions between Ubvs and other HECT-like E3 ligases (23). Interestingly, Ubv A06 and Ubv D09 do not possess any mutations in region 1 of the phage display library. Mutations in this region are common amongst Ubvs that were identified as high affinity binders of other classes of human enzymes, suggesting that SspH1 may not interact with this surface of ubiquitin (32).

Expression of either Ubv in *S. cerevisiae* did not lead to a detectable growth defect on solid media or in liquid media. Although it has been previously shown that expression of ubiquitin containing a mutation at the R74 residue has a dominant negative effect on yeast growth, our observation that yeast growth is not impacted despite the presence of this mutation may be attributed to the selective nature of Ubvs, as they are known to have high specificity for their cognate protein (53–55). However, it has also been reported that an intact C-terminal diglycine motif is required for the dominant negative effect of R74 to be observed (53). It is also notable that yeast ubiquitin differs from human ubiquitin at 3 residues (Ser 19, Asp 24, Ser 28), none of which are found within either Ubv (56). Consistent with previous findings, we observed SspH1- mediated toxicity in yeast dependent on the catalytic activity of SspH1 (13). Co-expression of Ubvs alongside SspH1 was sufficient to rescue yeast growth relative to the non-induced condition in liquid media. Conversely, Ubv co-expression on solid media only partially rescued yeast growth. These observations may be attributed to the different environmental pressures experienced by yeast growing in liquid or on solid media as well as the previously observed effects of ubiquitin overexpression (57).

Despite yeast toxicity being a known consequence of SspH1 expression in *S. cerevisiae* for over a decade, the mechanism behind this phenomenon is not fully understood. Here we report an increase in cell cycle perturbations, notably the inability for yeast to progress through the G2/M phase of the cell cycle, by both microscopic and cytometric analyses in the presence of SspH1. This cell cycle interference phenotype was dependent on the catalytic activity of SspH1 and was suppressed in the presence of either Ubv. Interestingly, the interaction between SspH1 and PKN1 was initially identified through a yeast two-hybrid screen suggesting the presence of a preferred substrate is sufficient to suppress SspH1-mediated toxicity (15,18). We also observed no detrimental effect of Ubv expression on yeast cell cycle progression, consistent with previous observations that they do not impact yeast growth (53).

Intriguingly, despite our hypothesis that Ubvs would interfere with SspH1 E3 ubiquitin ligase activity, based upon our yeast studies, we report an increase in recombinant SspH1 *in vitro* E3 ubiquitin ligase activity in the presence of Ubv D09 and human ubiquitin. Nevertheless, our studies with Ubv D09 revealed it could not be polymerized into polyubiquitin chains, as expected, given that Ubv D09 lacks the C-terminal diglycine motif necessary for the formation of thioester linkage (50,58). The slight reduction of SspH1 activity observed in the presence of ΔDiGly may also be owed to the lack of a C-terminal diglycine motif (58). Interestingly, ΔDiGly did not reduce SspH1 activity in the presence of PKN1, which may be attributed to the increase in activity NELs are known to undergo in the presence of their cognate substrate (41). By contrast, Ubv D09 enhanced SspH1 activity in the presence and absence of PKN1, although the linkage pattern of PKN1-ubiquitination was altered by Ubv D09. Lys48-linked chains are typically associated with proteasomal degradation and have recently been shown to be the primary polyubiquitin linkage formed by SspH1, which is consistent with its described role in mediating PKN1 degradation (18,20). Our observations confirm that SspH1-mediated ubiquitination primarily consists of Lys48-linked ubiquitin chains but that this composition can be modulated by a ubiquitin variant. It has been previously observed that the presence of Ubv can affect the natural bias of ubiquitin distribution of an E3 ligase, altering the ratio of processive and distributive ubiquitination of the substrate (34). Our experiments were limited to assessing K48- and K63-Ub linkages and we cannot rule out that Ubv D09 had an impact on other linkage types. Nevertheless, it is tempting to speculate that the basis of SspH1 toxicity in yeast is the formation of K48-ubiquitin linkages on, and subsequent degradation of, a yeast ortholog of PKN1. Expression of Ubv D09 in yeast may reduce K48 linkage on this unknown substrate, below a threshold that mitigates yeast toxicity. Despite the disconnect between the effects of Ubv D09 in the *S. cerevisiae* model and *in vitro*, we consistently observe a significant modulation of the SspH1 phenotype in both cases. It will be interesting to revisit *in vitro* ubiquitination assays of SspH1, Ubv D09 and the yeast substrate, once it is identified.

Although NELs are present amongst several well-studied Gram-negative bacterial species, their unique structure has limited the available tools to probe their molecular mechanisms (8). Given that Ubvs have been previously demonstrated to be highly selective between enzymes of the same family, they may also be employed to probe the level of redundancy that exists between the closely related effectors (8,55). To our knowledge, this is the first report that demonstrates an Ubv approach can be employed to identify modulators of a bacterial-encoded E3 ubiquitin ligase. Further studies are required to elucidate the biochemical interaction between the Ubvs and SspH1 which may provide information on the unique biology behind NEL effectors.

Additionally, the identification of additional Ubvs which are high-affinity binders to other NELs is required to discern the relative selectivity of this approach amongst the effector family.

Nevertheless, our current work indicates that a Ubv approach initially intended to target a human family of ubiquitin ligases can be successfully repurposed to target bacterial effectors with a unique, convergently evolved mechanism of action, which provides an additional tool to probe the functional and mechanistic attributes of these effectors.

## Acknowledgements

This work was supported by operating grants from the Natural Sciences and Engineering Research Council of Canada (RGPIN-2020-04359 to APB) and the Government of Alberta Major Innovation Fund for Antimicrobial Research- One Health Consortium (RCP-19-003-MIF to APB). This research has also been funded by the Li Ka Shing Institute of Virology (LKSIoV). BED was supported by studentships from the University of Alberta Faculty of Medicine & Dentistry and LKSIoV. APB holds a Canada Research Chair (Tier 2) in Pattern Recognition Receptor Pathophysiology and this research was undertaken, in part, thanks to funding from the 10 Canada Research Chairs Program (231622). Some experiments were performed at the University of Alberta Faculty of Medicine & Dentistry Flow Cytometry Facility, RRID:SCR_019195, which receives financial support from the Faculty of Medicine & Dentistry and Canada Foundation for Innovation (CFI) awards to contributing investigators.

## Data Availability Statement

Flow cytometry data is available on flowrepository.org under experiment ID: FR-FCM-Z7AZ.

## Author Contributions

Designed research: BED AW WZ AM GE SSS APB

Performed research: BED AW WZ AM ASS

Analyzed data: BED APB

Wrote the paper, or provide their own descriptions: BED APB

Secured Funding: SSS APB

---

**S1 Fig.**
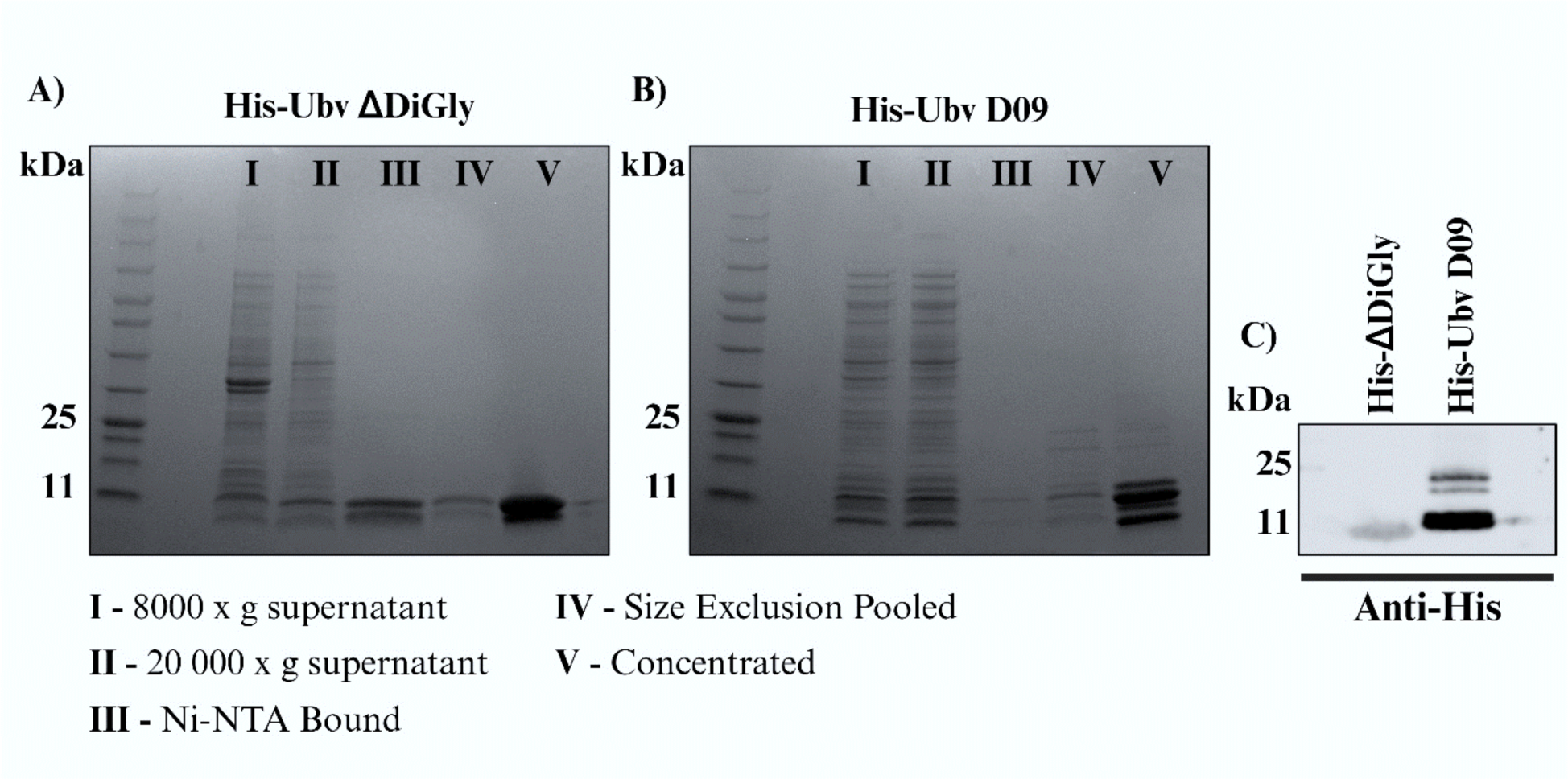
Purification of ΔDiGly and Ubv D09. **(A)** Purification of His_6_ – Ubv ΔDiGly ubiquitin. Lanes are as indicated. **(B)** Purification of His_6_ Ubv D09. Lanes are as indicated is panel A **(C)** Purified His_6_ – Ubv ΔDiGly ubiquitin and His_6_ –Ubv D09 were assessed using an anti-His antibody immunoblot.

**S2 Fig.**
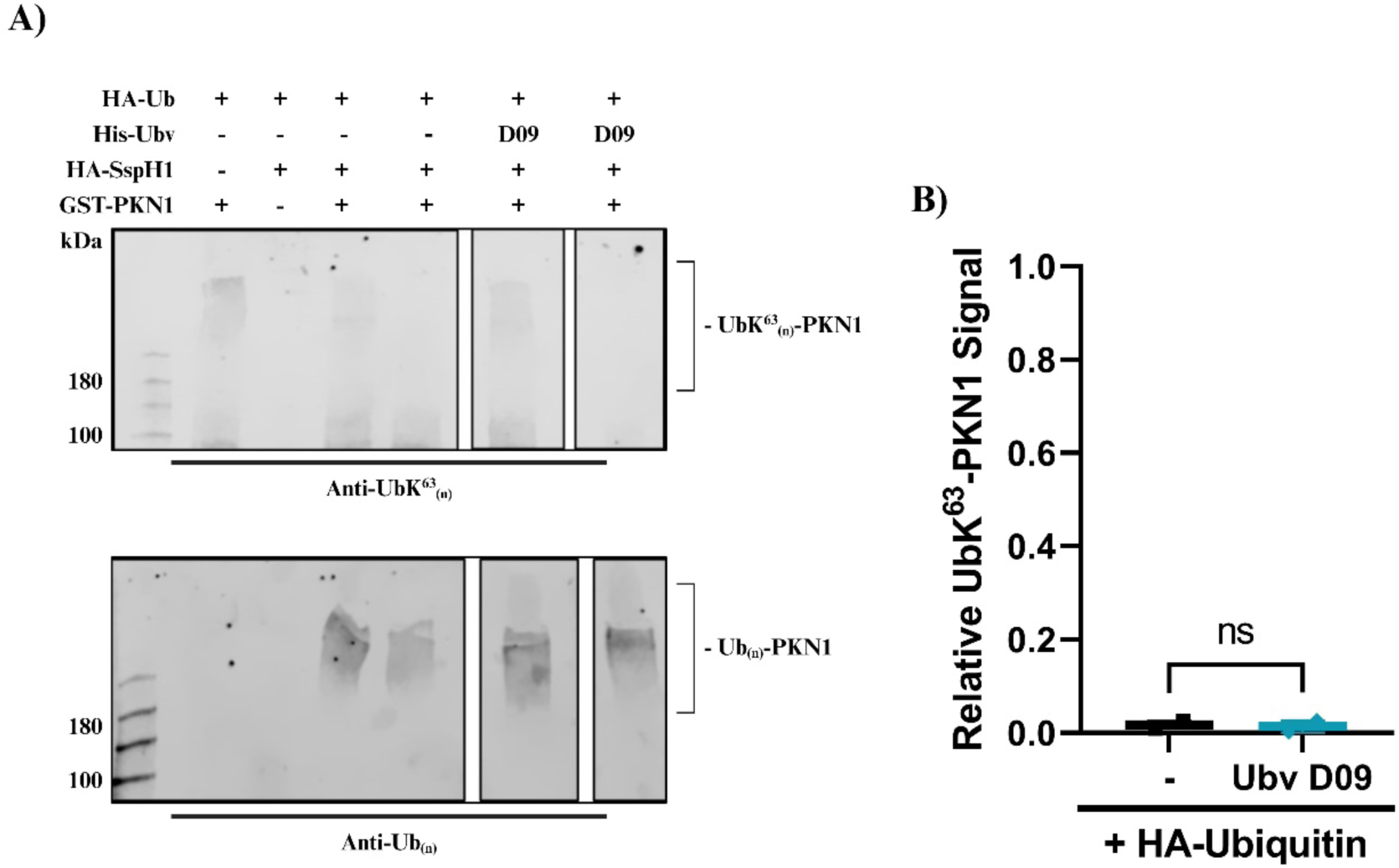
Ubvs modulate K63-linked SspH1-mediated ubiquitination of PKN1 *in vitro*. **(A)** Lysine-specific ubiquitin chain conformation of PKN1-specific, SspH1-mediated ubiquitination was determined by *in vitro* ubiquitination assays containing recombinant E1, E2, SspH1, PKN1, HA-Ub, or His-Ubv D09 as indicated and analyzed by immunoblot. Two independent reactions are shown. Total ubiquitination was determined by anti-Ub_(n)_ [P4D1], K63-specific ubiquitin chains was determined by anti-Ub^K63^ _(n)_[D7A11] **(B)** Formation of UbK^63^_(n)_-PKN1 and Ub_(n)_-PKN1 is expressed as ratio relative to the signal of Ub_(n)_-PKN1. Error bars represent the standard error of the mean across 4 independent experiments. Data was analyzed using an unpaired T-test.

**S3 Fig.**
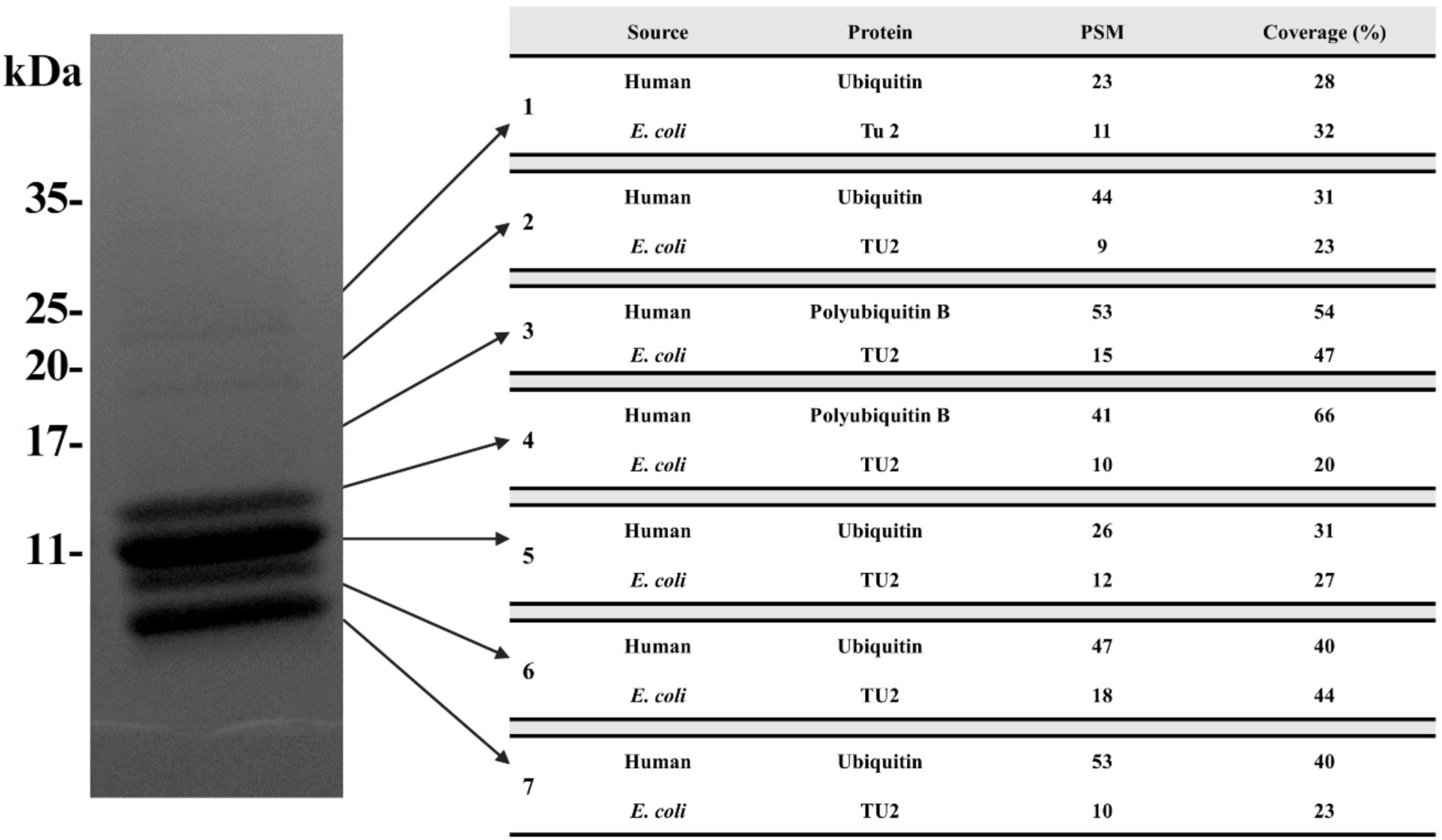
Mass Spectrometry of Purified Ubv D09. Purfied His-Ubv D09 was analyzed by SDS-PAGE on a 4-20% gradient gel. Bands were excised at the indicated locations and subjected to mass spectrometry using an Orbitrap Q Exactive mass spectrometer. Data was processed using Proteome Discoverer 1.4 and the Human and *E. coli* proteomic databases were searched using SEQUEST. The most abundant protein found in every excised band is shown alongside the most abundant contaminant. (PSM = Peptide Spectral Matches)

**S4 Fig.**
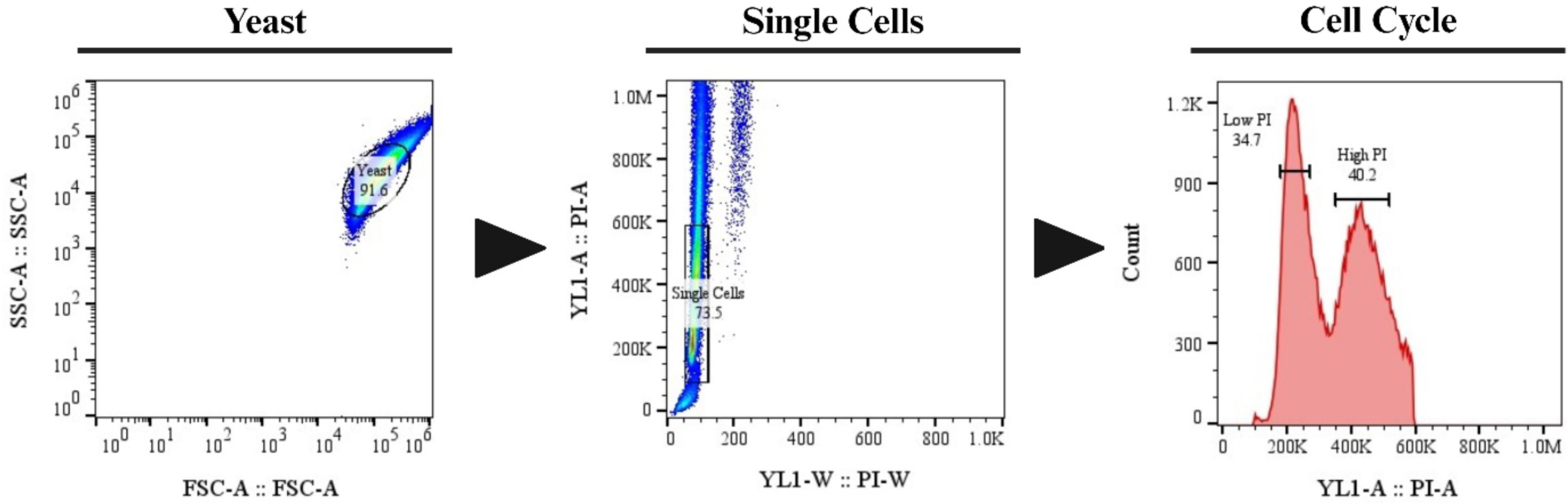
Gating Strategy for Flow Cytometry Analysis of Cell Cycle. Yeast were identified, and debris was excluded, using a forward scatter area (FSC-A) versus side scatter area (SSC-A) gate. Single cells were then selected on a YL1/PI-W versus YL1/PI-A plot to exclude doublets. Cell cycle analysis was then performed in this cell population by quantifying the ratio of cells with low PI, equivalent to 1N DNA content, and high PI, equivalent to 2N DNA content, fluorescent signal.

## References

1. Gal-Mor O, Boyle EC, Grassl GA. Same species, different diseases: how and why typhoidal and non-typhoidal Salmonella enterica serovars differ. Front Microbiol. 2014 Aug 4;5:102622.

2. Nemhauser J. Travel-Associated Infections & Diseases. In: Centers for Disease Control and Prevention (CDC), Nemhauser JB, editors. CDC Yellow Book 2024: Health Information for International Travel [Internet]. Oxford University Press; 2023 [cited 2024 Mar 19]. p. 0. Available from: 10.1093/oso/9780197570944.003.0005

3. Hume PJ, Singh V, Davidson AC, Koronakis V. Swiss Army Pathogen: The Salmonella Entry Toolkit. Front Cell Infect Microbiol [Internet]. 2017 [cited 2024 Feb 15];7. Available from: https://www.frontiersin.org/articles/10.3389/fcimb.2017.00348

4. Pillay TD, Hettiarachchi SU, Gan J, Diaz-Del-Olmo I, Yu XJ, Muench JH, et al. Speaking the host language: how Salmonella effector proteins manipulate the host. Microbiology. 2023;169(6):001342.

5. Galán JE. Salmonella Typhimurium and inflammation: a pathogen-centric affair. Nat Rev Microbiol. 2021 Nov;19(11):716–25.

6. Liu X, Jiang Z, Liu Z, Li D, Liu Z, Dong X, et al. Research Progress of Salmonella Pathogenicity Island. Int J Biol Life Sci. 2023 May 22;2(3):7–11.

7. Rohde JR, Breitkreutz A, Chenal A, Sansonetti PJ, Parsot C. Type III Secretion Effectors of the IpaH Family Are E3 Ubiquitin Ligases. Cell Host Microbe. 2007;1(1):77–83.

8. Bullones-Bolaños A, Bernal-Bayard J, Ramos-Morales F. The NEL Family of Bacterial E3 Ubiquitin Ligases. Int J Mol Sci. 2022 Jan;23(14):7725.

9. Norkowski S, Schmidt MA, Rüter C. The species-spanning family of LPX-motif harbouring effector proteins. Cell Microbiol. 2018;20(11):e12945.

10. Singer AU, Schulze S, Skarina T, Xu X, Cui H, Eschen-Lippold L, et al. A Pathogen Type III Effector with a Novel E3 Ubiquitin Ligase Architecture. PLOS Pathog. 2013 Jan 24;9(1):e1003121.

11. Iyer LM, Burroughs AM, Aravind L. The prokaryotic antecedents of the ubiquitin-signaling system and the early evolution of ubiquitin-like β-grasp domains. Genome Biol. 2006 Jul 19;7(7):R60.

12. Chou YC, Keszei AFA, Rohde JR, Tyers M, Sicheri F. Conserved Structural Mechanisms for Autoinhibition in IpaH Ubiquitin Ligases. J Biol Chem. 2012 Jan 2;287(1):268–75.

13. Keszei AFA, Tang X, McCormick C, Zeqiraj E, Rohde JR, Tyers M, et al. Structure of an SspH1-PKN1 Complex Reveals the Basis for Host Substrate Recognition and Mechanism of Activation for a Bacterial E3 Ubiquitin Ligase. Mol Cell Biol. 2013;34(3):362–73.

14. Miao EA, Scherer CA, Tsolis RM, Kingsley RA, Adams LG, Bäumler AJ, et al. Salmonella typhimurium leucine-rich repeat proteins are targeted to the SPI1 and SPI2 type III secretion systems. Mol Microbiol. 1999;34(4):850–64.

15. Haraga A, Miller SI. A Salmonella type III secretion effector interacts with the mammalian serine/threonine protein kinase PKN1. Cell Microbiol. 2006;8(5):837–46.

16. Haraga A, Miller SI. A Salmonella enterica Serovar Typhimurium Translocated Leucine-Rich Repeat Effector Protein Inhibits NF-κB-Dependent Gene Expression. Infect Immun. 2003 Jul;71(7):4052–8.

17. Bierne H, Pourpre R. Bacterial Factors Targeting the Nucleus: The Growing Family of Nucleomodulins. Toxins. 2020 Apr;12(4):220.

18. Bullones-Bolaños A, Martín-Muñoz P, Vallejo-Grijalba C, Bernal-Bayard J, Ramos-Morales F. Specificities and redundancies in the NEL family of bacterial E3 ubiquitin ligases of Salmonella enterica serovar Typhimurium. Front Immunol. 2024 Feb 1;15:1328707.

19. Metzger E, Müller JM, Ferrari S, Buettner R, Schüle R. A novel inducible transactivation domain in the androgen receptor: implications for PRK in prostate cancer. EMBO J. 2003 Jan 15;22(2):270–80.

20. Herod A, Emond-Rheault JG, Tamber S, Goodridge L, Lévesque RC, Rohde J. Genomic and phenotypic analysis of SspH1 identifies a new Salmonella effector, SspH3. Mol Microbiol [Internet]. 2021 [cited 2022 Jan 5]; Available from: http://onlinelibrary.wiley.com/doi/abs/10.1111/mmi.14871

21. Gül E, Huuskonen J, Abi Younes A, Maurer L, Enz U, Zimmermann J, et al. Salmonella T3SS-2 virulence enhances gut-luminal colonization by enabling chemotaxis-dependent exploitation of intestinal inflammation. Cell Rep. 2024 Mar 7;43(3):113925.

22. Middleton AJ, Teyra J, Zhu J, Sidhu SS, Day CL. Identification of Ubiquitin Variants That Inhibit the E2 Ubiquitin Conjugating Enzyme, Ube2k. ACS Chem Biol. 2021 Sep 17;16(9):1745–56.

23. LeBlanc N, Mallette E, Zhang W. Targeted modulation of E3 ligases using engineered ubiquitin variants. FEBS J. 2021;288(7):2143–65.

24. Zhang W, Bailey-Elkin BA, Knaap RCM, Khare B, Dalebout TJ, Johnson GG, et al. Potent and selective inhibition of pathogenic viruses by engineered ubiquitin variants. PLoS Pathog. 2017;13(5):e1006372.

25. McAlpine JM, Zhu J, Pudjihartono N, Teyra J, Currie MJ, Dobson RC, et al. Structural and biophysical characterisation of ubiquitin variants that specifically inhibit the ubiquitin conjugating enzyme Ube2d2 [Internet]. bioRxiv; 2024 [cited 2024 Mar 12]. p. 2024.03.10.583603. Available from: https://www.biorxiv.org/content/10.1101/2024.03.10.583603v1

26. Jansen G, Wu C, Schade B, Thomas DY, Whiteway M. Drag &Drop cloning in yeast. Gene. 2005 Jan 3;344:43–51.

27. Hartley JL, Temple GF, Brasch MA. DNA cloning using in vitro site-specific recombination. Genome Res. 2000 Nov;10(11):1788–95.

28. Raran-Kurussi S, Waugh DS. Expression and Purification of Recombinant Proteins in Escherichia coli with a His6 or Dual His6-MBP Tag. Protein Crystallogr. 2017 Jun 2;1607:1– 15.

29. Pettersen EF, Goddard TD, Huang CC, Couch GS, Greenblatt DM, Meng EC, et al. UCSF Chimera--a visualization system for exploratory research and analysis. J Comput Chem. 2004 Oct;25(13):1605–12.

30. Bhavsar AP, Brown NF, Stoepel J, Wiermer M, Martin DDO, Hsu KJ, et al. The Salmonella Type III Effector SspH2 Specifically Exploits the NLR Co-chaperone Activity of SGT1 to Subvert Immunity. PLOS Pathog. 2013;9(7):e1003518.

31. Lauman P, Dennis JJ. Synergistic Interactions among Burkholderia cepacia Complex- Targeting Phages Reveal a Novel Therapeutic Role for Lysogenization-Capable Phages. Microbiol Spectr [Internet]. 2023 Jun [cited 2023 Sep 18];11(3). Available from: https://www.ncbi.nlm.nih.gov/pmc/articles/PMC10269493/

32. Leung I, Jarvik N, Sidhu SS. A Highly Diverse and Functional Naïve Ubiquitin Variant Library for Generation of Intracellular Affinity Reagents. J Mol Biol. 2017;429(1):115–27.

33. Ernst A, Avvakumov G, Tong J, Fan Y, Zhao Y, Alberts P, et al. A Strategy for Modulation of Enzymes in the Ubiquitin System. Science. 2013 Feb;339(6119):590–5.

34. Zhang W, Wu KP, Sartori MA, Kamadurai HB, Ordureau A, Jiang C, et al. System-Wide Modulation of HECT E3 Ligases with Selective Ubiquitin Variant Probes. Mol Cell. 2016 Apr 7;62(1):121–36.

35. Waterhouse A, Bertoni M, Bienert S, Studer G, Tauriello G, Gumienny R, et al. SWISS- MODEL: homology modelling of protein structures and complexes. Nucleic Acids Res. 2018;46(Journal Article):W296–303.

36. Pickart CM, Eddins MJ. Ubiquitin: structures, functions, mechanisms. Biochim Biophys Acta BBA - Mol Cell Res. 2004 Nov 29;1695(1):55–72.

37. Evans R, O’Neill M, Pritzel A, Antropova N, Senior A, Green T, et al. Protein complex prediction with AlphaFold-Multimer [Internet]. bioRxiv; 2022 [cited 2023 Sep 19]. p. 2021.10.04.463034. Available from: https://www.biorxiv.org/content/10.1101/2021.10.04.463034v2

38. Popa C, Coll NS, Valls M, Sessa G. Yeast as a Heterologous Model System to Uncover Type III Effector Function. PLoS Pathog. 2016 Feb 25;12(2):e1005360.

39. Slagowski NL, Kramer RW, Morrison MF, LaBaer J, Lesser CF. A Functional Genomic Yeast Screen to Identify Pathogenic Bacterial Proteins. PLOS Pathog. 2008 Jan 18;4(1):e9.

40. Sloper-Mould KE, Jemc JC, Pickart CM, Hicke L. Distinct Functional Surface Regions on Ubiquitin *. J Biol Chem. 2001 Aug 10;276(32):30483–9.

41. Quezada CM, Hicks SW, Galán JE, Stebbins CE, Bassler BL. A Family of Salmonella Virulence Factors Functions as a Distinct Class of Autoregulated E3 Ubiquitin Ligases. Proc Natl Acad Sci - PNAS. 2009;106(12):4864–9.

42. Singer AU, Rohde JR, Lam R, Skarina T, Kagan O, DiLeo R, et al. Structure of the Shigella T3SS effector IpaH defines a new class of E3 ubiquitin ligases. Nat Struct Mol Biol. 2008 Dec;15(12):1293–301.

43. Mager WH, Winderickx J. Yeast as a model for medical and medicinal research. Trends Pharmacol Sci Regul Ed. 2005;26(5):265–73.

44. Calvert MEK, Lannigan JA, Pemberton LF. Optimization of yeast cell cycle analysis and morphological characterization by multispectral imaging flow cytometry. Cytometry A. 2008;73A(9):825–33.

45. Chiou J geng, Balasubramanian MK, Lew DJ. Cell Polarity in Yeast. Annu Rev Cell Dev Biol. 2017 Oct 6;33(1):77–101.

46. Foltman M, Molist I, Sanchez-Diaz A. Synchronization of the Budding Yeast Saccharomyces cerevisiae. In: Sanchez-Diaz A, Perez P, editors. Yeast Cytokinesis: Methods and Protocols [Internet]. New York, NY: Springer; 2016 [cited 2021 Jul 16]. p. 279–91. (Methods in Molecular Biology). Available from: 10.1007/978-1-4939-3145-3_19

47. Greenwood BL, Stuart DT. Synchronization of Saccharomyces cerevisiae Cells for Analysis of Progression Through the Cell Cycle. In: Wang Z, editor. Cell-Cycle Synchronization: Methods and Protocols [Internet]. New York, NY: Springer US; 2022 [cited 2023 Dec 13]. p. 145–68. (Methods in Molecular Biology). Available from: 10.1007/978-1-0716-2736-5_12

48. Künkel W. Effects of the antimicrotubular cancerostatic drug nocodazole on the yeast Saccharomyces cerevisiae. Z Für Allg Mikrobiol. 1980;20(5):315–24.

49. Cook M, Delbecq SP, Schweppe TP, Guttman M, Klevit RE, Brzovic PS. The ubiquitin ligase SspH1 from Salmonella uses a modular and dynamic E3 domain to catalyze substrate ubiquitylation. J Biol Chem. 2018 Nov 20;294(3):783–93.

50. Zhao B, Bhuripanyo K, Schneider J, Zhang K, Schindelin H, Boone D, et al. Specificity of the E1-E2-E3 Enzymatic Cascade for Ubiquitin C-Terminal Sequences Identified by Phage Display. ACS Chem Biol. 2012 Dec 21;7(12):2027–35.

51. Newton K, Matsumoto ML, Wertz IE, Kirkpatrick DS, Lill JR, Tan J, et al. Ubiquitin Chain Editing Revealed by Polyubiquitin Linkage-Specific Antibodies. Cell. 2008 Aug 22;134(4):668–78.

52. Franklin TG, Pruneda JN. Bacteria make surgical strikes on host ubiquitin signaling. PLOS Pathog. 2021 Mar 18;17(3):e1009341.

53. Padhy AA, Mavor D, Sahoo S, Bolon DNA, Mishra P. Systematic profiling of dominant ubiquitin variants reveals key functional nodes contributing to evolutionary selection. Cell Rep. 2023 Sep 26;42(9):113064.

54. Duerksen-Hughes PJ, Xu X, Wilkinson KD. Structure and function of ubiquitin: evidence for differential interactions of arginine-74 with the activating enzyme and the proteases of ATP-dependent proteolysis. Biochemistry. 1987 Nov 1;26(22):6980–7.

55. Veggiani G, Yates BP, Martyn GD, Manczyk N, Singer AU, Kurinov I, et al. Panel of Engineered Ubiquitin Variants Targeting the Family of Human Ubiquitin Interacting Motifs. ACS Chem Biol. 2022 Apr 15;17(4):941–56.

56. Vijay-Kumar S, Bugg CE, Wilkinson KD, Vierstra RD, Hatfield PM, Cook WJ. Comparison of the three-dimensional structures of human, yeast, and oat ubiquitin. J Biol Chem. 1987 May 5;262(13):6396–9.

57. Chen Y, Piper PW. Consequences of the overexpression of ubiquitin in yeast: elevated tolerances of osmostress, ethanol and canavanine, yet reduced tolerances of cadmium, arsenite and paromomycin. Biochim Biophys Acta BBA - Mol Cell Res. 1995 Jul 20;1268(1):59–64.

58. Pickart CM, Kasperek EM, Beal R, Kim A. Substrate properties of site-specific mutant ubiquitin protein (G76A) reveal unexpected mechanistic features of ubiquitin-activating enzyme (E1). J Biol Chem. 1994 Mar 11;269(10):7115–23.

59. Auweter SD, Bhavsar AP, de Hoog CL, Li Y, Chan YA, van der Heijden J, et al. Quantitative Mass Spectrometry Catalogues Salmonella Pathogenicity Island-2 Effectors and Identifies Their Cognate Host Binding Partners. J Biol Chem. 2011 Jul 8;286(27):24023–35.

